# An Rhs effector uses distinct target cell functions to intoxicate bacterial and fungal competitors

**DOI:** 10.1101/2025.10.28.685041

**Authors:** Gabriela Mol Avelar, Genady Pankov, Tina Sarapa, Janet Quinn, William N. Hunter, Colin Rickman, Sarah J. Coulthurst

## Abstract

Many bacteria use Type VI secretion systems (T6SSs) to deliver toxic effector proteins into neighbouring bacterial or fungal cells as a means of inter-microbial competition. Compared with numerous antibacterial effectors, few antifungal effectors have been described. Furthermore, how T6SS-delivered effectors reach their site of action in different types of target cell remains poorly understood. Here, we combine structural biology with in vivo approaches to show that Rhs2 from *Serratia marcescens* Db10 is a dual-kingdom T6SS-dependent DNase effector which hijacks distinct, essential target cell functions in order to reach its site of action in bacterial and fungal cells. In bacterial cells, the Rhs2 toxin domain (Rhs2CT) interacts specifically with the elongation factor, EF-Tu, and, in sibling cells, interacts with the cognate immunity protein in an unusual manner. Interaction with EF-Tu is essential for T6SS-mediated intoxication of bacterial cells by Rhs2, but not for DNase activity or intoxication of fungal cells, implying it facilitates entry of Rhs2CT across the inner membrane to the cytoplasm. Alternatively, in fungal cells, Rhs2CT translocates to the nucleus using the nuclear import machinery. Our findings reveal how a single effector domain can act against targets with distinct cellular architectures and suggest that dual-kingdom effectors may occur widely.

## INTRODUCTION

Microorganisms typically exist in polymicrobial communities whose members interact co-operatively and competitively to maximise their access to resources and space. Bacteria employ a variety of mechanisms to actively compete with each other and with other microbes, including production of diffusible antimicrobial molecules and sophisticated machineries for contact-dependent delivery of protein toxins^1, 2^. Such competitive interactions are critical in determining the composition and dynamics of diverse polymicrobial communities, with roles in promoting incursion by pathogenic bacteria, microbiota-mediated defence against incoming colonisers, establishment of symbiotic and environmental communities, and promoting microbial evolution^1, 2^. A widespread mechanism for inter-bacterial competition is the Type VI secretion system (T6SS), a nanoweapon used to deliver multiple toxic effector proteins into neighbouring cells^3^. A large number of antibacterial effector proteins delivered by the T6SS have been identified, including several classes of peptidoglycan hydrolases, phospholipases, DNA nucleases and deaminases, NADases, ADP-ribosyltransferases and pore-forming toxins^4, 5^. In each case, the secreting cell encodes a specific immunity protein able to provide protection against the action of the cognate effector, including incoming effectors delivered by genetically identical sibling neighbours. More recently, it has been shown that the T6SS can also be used against microbial fungi, extending the breadth of its ability to influence polymicrobial communities and suggesting potential future therapeutic or agricultural applications against fungal pathogens^6, 7^. However, to date, only a handful of anti-fungal effector proteins have been reported.

The basic mechanism of effector delivery by the T6SS is well understood^5, 8, 9^. Contraction of an extended cytoplasmic tubular sheath structure, anchored on a membrane-bound basal complex, propels an arrow-like puncturing structure, decorated with effector proteins, out of the secreting cell and into a neighbouring target cell. The expelled puncturing structure comprises a tube of stacked hexameric rings of Hcp protein, topped with a spike complex made of a trimer of VgrG proteins and a single PAAR domain-containing protein forming the sharp tip of the spike. Effectors associate with this structure for delivery into a target cell in several ways^4, 8^: Cargo effectors interact non-covalently with Hcp, VgrG or PAAR proteins, whilst specialised effectors comprise an effector domain covalently fused to the C-terminal domain of an Hcp, VgrG or PAAR protein. Hcp-dependent effectors sit in the lumen of the Hcp tube whilst VgrG- and PAAR-dependent effectors are located on the outside of the VgrG-PAAR spike. In some cases, additional adaptor or chaperone proteins are required to stabilise the effector and/or recruit it to the puncturing structure^5, 8^. PAAR domain-containing Rhs (Rearrangement hot spot) proteins are large polymorphic toxins which represent a widespread class of specialised T6SS effectors^10^. The N-terminal region of these Rhs proteins contains the PAAR domain forming the tip of the T6SS puncturing structure and a transmembrane domain involved in translocation of the effector domain to the cytoplasm of recipient cells^11^. The middle of the protein contains Rhs repeats forming a β-barrel cage-like structure around the C-terminal toxin domain (CT) and terminates with a highly conserved region including an autoproteolytic cleavage site for release of the CT^12, 13^. Rhs CTs are highly variable, even within species, and can be readily exchanged as a functional unit with the cognate immunity protein encoded immediately downstream by homologous recombination^14, 15^. In addition to their effector function, which can be very potent, these Rhs proteins also have a structural function within the T6SS, interacting with a specific VgrG protein to provide the essential PAAR domain tip for assembly and firing of the expelled puncturing structure^16^. Compared with T6SS assembly and effector loading in the secreting cell, the fate of the effector in the recipient cell is much less well understood, including the mechanisms by which effectors and effector domains which act in the cytoplasm reach this compartment. Whilst recent work has provided a few specific examples to show that T6SS effectors can contain functions to aid them in crossing the bacterial inner membrane and can hijack host proteins to assist their toxicity^11, 17, 18^, how T6SS effectors reach their final destination in an active state remains a key question for the field.

*Serratia marcescens* is an opportunistic pathogen which is found widely in the environment but also represents an important cause of hospital-acquired infections, including those complicated by antibiotic resistance^19^. We have used the model strain Db10 to show that a conserved T6SS encoded by all *S. marcescens* and much of the *Serratia* genus, has potent antibacterial and antifungal activity^7, 20, 21^. *S. marcescens* Db10 secretes at least eight antibacterial T6SS effectors, including two peptidoglycan amidases, an NADase and an ion-selective pore-forming toxin^22, 23, 24^. The Rhs2 protein is a PAAR-containing effector and potent anti-bacterial toxin whose CT was indirectly inferred to have nuclease activity^25^. Rhs2 interacts with one of the two VgrG proteins in Db10, VgrG2, to form the most efficient version of the Db10 T6SS, although two alternative spike complexes (VgrG1-Paar1, VgrG2-Rhs1) can also support function^16^. The T6SS of *S. marcescens* Db10 also delivers two antifungal effectors, Tfe1 and Tfe2, whose action is specific to fungal cells^7^. However, in this previous study, we noted that Tfe1 and Tfe2 cannot account for all the anti-fungal activity of the *S. marcescens* Db10 T6SS, since a Δ*tfe1*Δ*tfe2* mutant displayed some antifungal activity compared with a T6SS inactive mutant, and, therefore, there must be at least one other effector with antifungal activity.

Here, we show that Rhs2 represents the remaining antifungal T6SS effector in *S. marcescens* Db10 and that it is a dual-kingdom effector whose C-terminal DNase domain hijacks distinct target cell functions in order to reach its site of action in bacterial and fungal cells. We show that the Rhs2 CT interacts specifically with bacterial Elongation Factor Thermo-unstable (EF-Tu) and that this interaction is essential for T6SS-mediated intoxication of bacterial cells but not for DNase activity or intoxication of fungal cells, implying it facilitates entry of the toxin across the inner membrane to the cytoplasm. Alternatively, in fungal cells, the Rhs2 CT translocates from the cytoplasm to the nucleus using the nuclear import machinery and nuclear DNA damage response mechanisms may provide some protection against toxicity. Overall, this study reveals how single effector domains can possess the versatility to act against targets with distinct cellular architectures.

## RESULTS

### Rhs2 is a dual-kingdom effector which acts against bacterial and fungal cells and represents the remaining antifungal effector delivered by the Type VI secretion system of *S. marcescens* Db10

In order to identify the effector responsible for the remaining T6SS-dependent antifungal activity observed in the absence of the fungal-specific effectors Tfe1 and Tfe2^7^, we considered the other T6SS effector proteins of *S. marcescens* Db10 identified previously. Initial work suggested that the C-terminal effector domain of Rhs2 (Rhs2CT) could be responsible. However, given the important structural role played by the PAAR domain-containing Rhs2 protein in overall T6SS function^16^, it was necessary to generate a strain of Db10 in which the toxicity of the C-terminal effector domain was eliminated, whilst the ability of the Rhs protein to support T6SS firing and delivery of other effectors was unaffected. Therefore, using the sequence similarity between Rhs2CT and other HNH endonucleases, we generated a strain encoding Rhs2 with a single amino acid substitution in a residue predicted to be essential for catalytic activity, H1369A (Rhs2_CM_, Rhs2 <u>c</u>atalytic <u>m</u>utant), at the normal chromosomal location. This strain, Db10 *rhs2_CM_*, was then tested for its ability to intoxicate a non-immune bacterial strain, Db10 Δ*rhs2*Δ*rhsI2*. The Δ*rhs2*Δ*rhsI2* mutant lacks the RhsI2 immunity protein which confers protection against Rhs2CT and is sensitive to the action of Rhs2 but not any of the other effectors delivered by Db10. Co-culture of wild type Db10 with the Rhs2-sensitive target strain led to a six-log drop in the recovery of viable target cells, emphasizing the potent anti-bacterial activity of Rhs2 (Figure 1A). In contrast, the *rhs2_CM_* mutant showed a complete loss of activity against this target, being indistinguishable from a full deletion of *rhs2* (Δ*rhs2*) or a T6SS-inactive mutant (Δ*tssE*). To confirm that the Rhs2_CM_ variant had no loss of its ability to support overall T6SS function, we used *Escherichia coli* as the target strain in co-culture assays for T6SS-dependent anti-bacterial activity. In this case, the target is killed by a combination of eight antibacterial effectors which can be delivered using Rhs2. In an otherwise wild type background, the *rhs2_CM_* mutant retained the ability to inhibit *E. coli* to a similar extent as the wild type, indicating that all the other antibacterial effectors are being delivered normally. However, the Δ*rhs2* mutant showed a reduction in overall T6SS-dependent antibacterial activity, confirming the previously-observed requirement for an intact PAAR-containing Rhs2 protein for full T6SS activity (Figure 1B)^16^. In a Δ*vgrG1*Δ*rhs1* background, where all T6SSs must use Rhs2 and therefore T6SS activity is entirely dependent on a functional Rhs2^16^, the *rhs2_CM_* mutant again showed no loss of activity compared with native *rhs2*, in contrast with the total loss of T6SS activity for the Δ*rhs2* deletion (Figure 1B). These data together confirm that the H1369A mutation in Rhs2_CM_ removes the effector function (toxicity) of Rhs2CT without compromising the structural function of full-length Rhs2.

**Figure 1.**
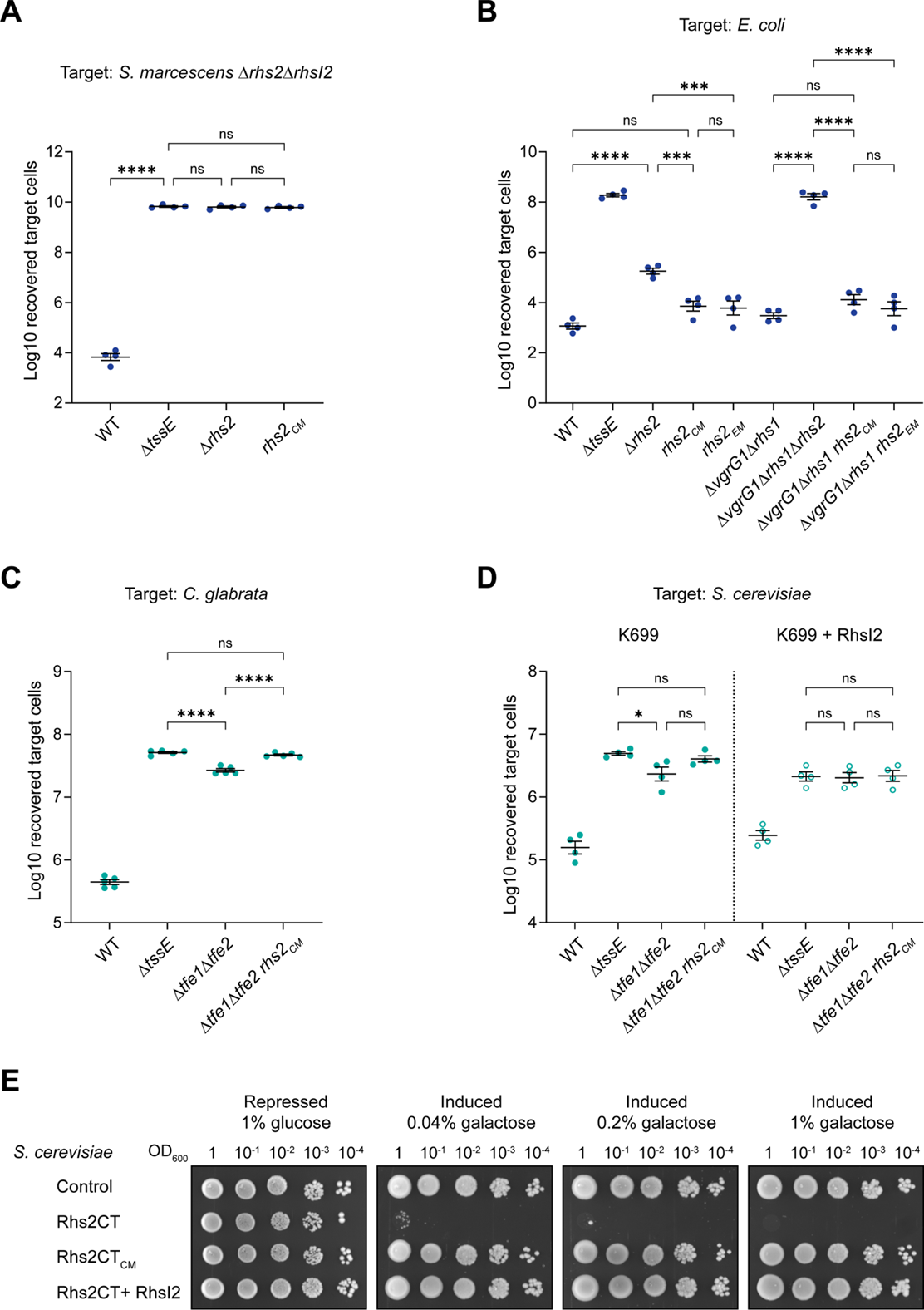
Rhs2 acts against bacterial and fungal cells and represents the remaining antifungal effector protein deployed by the T6SS of *S. marcescens* Db10 in addition to Tfe1 and Tfe2. **(A, B)** Recovery of target bacteria, either the Rhs2-susceptible Δ*rhs2*Δ*rhsI2* mutant of *S. marcescens* Db10 (A) or *E. coli* BW25113 (B), following co-culture with wild type *S. marcescens* Db10 (WT) or mutants carrying in-frame gene deletions or point mutations in *rhs2* as indicated (Rhs2_CM_, Rhs2 H1369A; Rhs2_EM_, Rhs2 E1341R R1344A A1378W K1380E). Data are presented as mean ± SEM with individual data points overlaid (n=4 biological replicates; **** P<0.0001, *** P<0.001, ns not significant; one-way ANOVA with Tukey’s test; for clarity, only selected comparisons are displayed). **(C, D)** Recovery of target fungal cells following co-culture with wild type Db10 or mutants carrying in-frame gene deletions or point mutations in *rhs2* as indicated. The fungal target is *C. glabrata* ATCC2001 in panel C, and *S. cerevisiae* K699 with a chromosomal integration construct directing the expression of RhsI2 (K699 + RhsI2) or the empty promoter control (K699) in panel D. Data are presented as mean ± SEM with individual data points overlaid (n=5 or n=4 biological replicates for panels C and D, respectively; **** P<0.0001, * P<0.05, ns not significant; one-way ANOVA with Šídák’s test). **(E)** Growth of *S. cerevisiae* K699 chromosomal integration strains carrying the empty promoter construct (control) or constructs directing the expression of the wild type Rhs2 C-terminal domain (Rhs2CT), the catalytically inactive variant (Rhs2CT_CM_, H1369A), or Rhs2CT with the immunity protein RhsI2 (Rhs2CT + RhsI2) on media containing glucose or galactose for repression or induction, respectively, of gene expression.

We then used the *rhs2_CM_* mutant to investigate the contribution of the effector function of Rhs2 to the T6SS-dependent antifungal activity of *S. marcescens* Db10. When Db10 was co-cultured with *Candida glabrata*, some antifungal activity remained in the absence of Tfe1 and Tfe2, as shown by a lower recovery of yeast cells with the Δ*tfe1*Δ*tfe2* mutant than the Δ*tssE* mutant. When the effector function of Rhs2 was also removed in the Δ*tfe1*Δ*tfe2* background (Δ*tfe1*Δ*tfe2 rhs2_CM_*), the remaining T6SS-dependent anti-fungal activity was lost (Figure 1C), demonstrating that Rhs2CT is also an antifungal effector and contributes the remaining antifungal activity of Db10 towards *C. glabrata*. Similar observations were made with *Saccharomyces cerevisiae* K699 as the target, where the Δ*tfe1*Δ*tfe2* mutant but not the Δ*tfe1*Δ*tfe2 rhs2_CM_* mutant showed reduced fungal recovery compared with the Δ*tssE* mutant (Figure 1D). Additionally, when the immunity protein, RhsI2, was expressed in K699, there was no longer a difference in recovery of the fungal cells between co-culture with the Δ*tfe1*Δ*tfe2* or the Δ*tssE* mutant, confirming that the residual Tfe1/Tfe2-independent antifungal activity of *S. marcescens* results from the delivery of Rhs2CT (Figure 1D). Finally, we confirmed directly that Rhs2CT is an antifungal toxin by heterologous expression in *S. cerevisiae* cells. Expression of wild type Rhs2CT was highly toxic, preventing yeast growth even at low induction. This toxicity could be prevented either by expressing the inactive variant (Rhs2_CM_) or by co-expressing the immunity protein RhsI2 (Figure 1E). Overall, these data demonstrate that Rhs2CT is a dual-kingdom effector which can be deployed in a T6SS-dependent manner against both bacterial and fungal cells.

### Rhs2CT is an atypical HNH endonuclease-type DNase which interacts with RhsI2 and also forms a heterotrimeric complex including EF-Tu

In previous work, we observed plasmid DNA degradation upon production of Rhs2CT in *E. coli* and noted that the sequence of Rhs2CT included a putative partial HNH endonuclease domain, indicating that this effector was likely to have nuclease activity^25^. To investigate the molecular function of Rhs2CT and to understand how its activity is neutralised by its cognate immunity protein, RhsI2, structural studies were initiated. His_6_-tagged Rhs2CT was co-produced with RhsI2 in *E. coli* in order to prevent toxicity and allow purification of the effector-immunity complex. Unexpectedly, the Rhs2CT-RhsI2 complex was found to co-purify with *E. coli* EF-Tu, a highly conserved and abundant protein required for protein synthesis. The complex with EF-Tu was purified to homogeneity and size-exclusion chromatography indicated that the three proteins form a stable 1:1:1 complex (Figure 2A).

**Figure 2.**
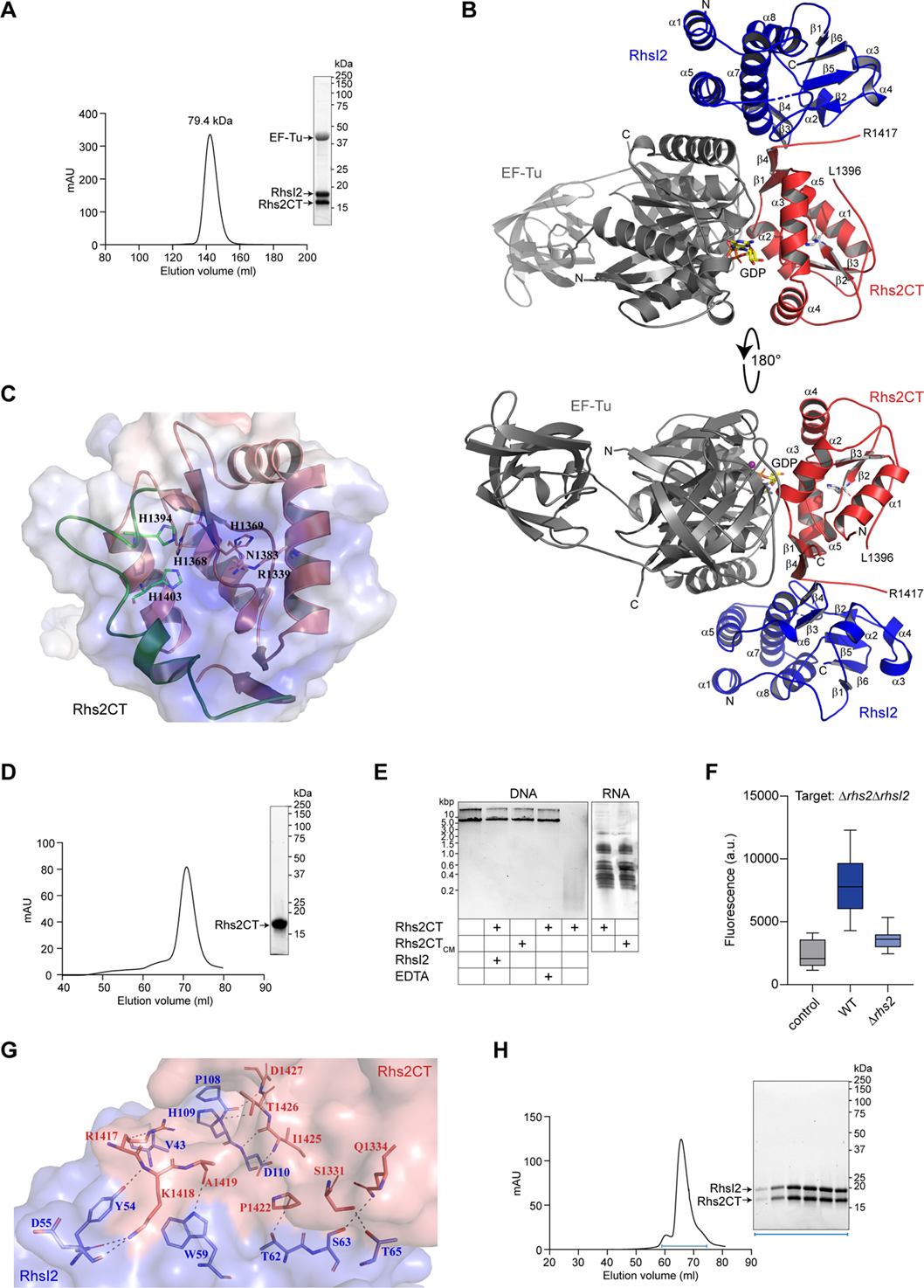
Rhs2CT is an atypical HNH endonuclease-type DNase which forms a heterotrimeric complex with RhsI2 and EF-Tu. **(A)** Size-exclusion chromatography using a Superdex 75 26/60 column of the protein complex isolated from *E. coli* producing recombinant His_6_-Rhs2CT and untagged RhsI2. Proteins from the peak were visualised by SDS-PAGE and Coomassie staining (inset) and the estimated molecular weight corresponding to the peak elution volume is noted. The protein copurifying with Rhs2CT-RhsI2 was identified as *E. coli* EF-Tu by mass spectrometry. **(B)** Overall arrangement of the Rhs2CT-RhsI2-EF-Tu complex, showing two views of the heterotrimer rotated by 180 degrees. The effector domain (Rhs2CT), immunity protein (RhsI2) and EF-Tu are coloured in red, blue and grey, respectively. GDP is shown as a stick and Mg^2+^ as a purple sphere. **(C)** Close-up depiction of the active site of Rhs2CT based on integration of crystallographic and AlphaFold models, with key residues shown as sticks and elements derived from AlphaFold shown in green. **(D)** Size-exclusion chromatography of refolded His_6_-Rhs2CT using a Superdex 75 16/60 column. **(E)** Nuclease activity of purified wild type Rhs2CT or catalytically-inactive Rhs2CT (Rhs2CT_CM_, H1369A) against plasmid DNA (left) or total bacterial RNA (right) in the presence or absence of RhsI2 or 1mM EDTA, as indicated. **(F)** TUNEL assay to detect DNA breaks in the Rhs2-susceptible Δ*rhs2*Δ*rhsI2* mutant of *S. marcescens* Db10 when co-cultured with wild type Db10 (WT), the Δ*rhs2* mutant, or no competitor (control). **(G)** Close-up depiction of interactions between effector (Rhs2CT, red) and immunity (RhsI2, blue) proteins. Residues of interest are shown as sticks and hydrogen bonds (within 2.5 Å-3.5 Å distance) are shown as dashed lines. **(H)** Size-exclusion chromatography using a Superdex 75 16/60 column of Rhs2CT-RhsI2 complex formed upon incubation of refolded His_6_-Rhs2CT and His_6_-RhsI2 at a 1:1 molar ratio. Fractions analysed by SDS-PAGE (inset) are indicated by the blue bracket.

The Rhs2CT-RhsI2-EF-Tu complex was crystallised and a 2.45 Å structure determined using molecular replacement (Figure 2B, Supplementary Figure 1). Search models included the structure of EF-Tu (PDB 1EFC), a preliminary structure of RhsI2, and a structure of Rhs2CT predicted by AlphaFold2^26^. The complex crystallised with two heterotrimers per asymmetric unit; however, the two heterotrimers appear to interact through crystal packing contacts between EF-Tu domain II which are unlikely to be of physiological significance. Both heterotrimers in the asymmetric unit are essentially identical and therefore the heterotrimer with the lowest *B*-factors was used for structural analysis. The Rhs2CT-RhsI2-EF-Tu complex forms a globular entity of approximately 85 × 65 × 55 Å, with a solvent-accessible surface area of 28,690 Å^2^. Within the complex, RhsI2 interacts with Rhs2CT via an extensive protein-protein interface, while Rhs2CT engages with the GDP-binding domain of EF-Tu (Figure 2B). The EF-Tu structure is fully resolved in the electron density; however, Rhs2CT is missing a 20-residue segment (Pro1397-Ser1416) that forms part of the active site, and RhsI2 lacks a four-residue segment (Lys74-Leu77) in a flexible loop region.

Rhs2CT adopts an α/β fold, comprising a central β-hairpin surrounded by α-helices, forming a compact and stable globular structure. The central β-hairpin (β2–β3) provides a core structural scaffold. Within β2, two histidines (His1368 and His1369) are predicted to play a key role in endonuclease activity (Figure 2C). His1369 forms part of the conserved HNH endonuclease motif, while the side chain of His1368 likely contributes to metal ion coordination within the active site. The polypeptide chain is not resolved between Leu1396 and Arg1417, where flexible loops link the structured regions. The missing segment, predicted to be part of the active site, is glycine- and proline-rich, suggesting that it represents a region of intrinsic flexibility or disorder, potentially involved in substrate binding or conformational dynamics (Supplementary Figure 1). Given this missing region, crystallographic and AlphaFold models of Rhs2CT were integrated in order to gain further insight into its active site and catalytic mechanism (Figure 2C). This analysis suggests that, as originally predicted, Rhs2CT belongs to the HNH endonuclease family (Pfam PF01844), with its active site resembling that of colicin E9 (PDB 1FSJ). However, Rhs2CT is an atypical member of the family, with its active site lacking the canonical ββα-metal fold, in which two residues involved in metal ion coordination are typically positioned on the α-helix of the ββα fold^27^. Instead, His1394 and His1403, located on the loop between β3 and α6, are predicted to form a metal-binding site together with His1368 on β2 (Figure 2C).

In order to confirm that Rhs2CT displays nuclease activity *in vitro*, it was isolated from the Rhs2CT-RhsI2 complex by an unfolding and refolding procedure (Figure 2D). Similar to colicin E9, Rhs2CT was able to degrade DNA in the presence of divalent cations such as Zn²⁺. Addition of the metal chelator EDTA, or of purified RhsI2, completely abolished nuclease activity, confirming that Rhs2CT is a metal-dependent nuclease (Figure 2E). Rhs2CT appears to be exclusively a DNase, with no detectable activity against RNA (Figure 2E). Finally, we used a TUNEL (terminal deoxynucleotidyl transferase dUTP nick end labeling) assay to confirm the presence of Rhs2-induced DNA breaks in intoxicated cells when Rhs2 is delivered into susceptible target cells by the T6SS (Figure 2F).

The structure of the immunity protein, RhsI2, is predominantly α-helical, with a single two-stranded antiparallel β-sheet at its core (Supplementary Figure 1) and represents a previously-undescribed structure with no homologues in the PDB. Unusually, compared with other T6SS-associated effector-immunity complexes^5^, RhsI2 does not inactivate Rhs2CT through direct occlusion of the active site. Instead, RhsI2 binds away from the active site (Figure 2B), suggesting an alternative inhibitory mechanism. This mechanism may involve RhsI2 inducing a conformational rearrangement in the loop spanning His1388-Gly1407 of Rhs2CT, which contains residues required for metal co-ordination (His1394 and His1403), since this loop appears partially disordered in both the heterotrimeric Rhs2CT-RhsI2-EF-Tu complex structure and a structure of the heterodimeric Rhs2CT-RhsI2 complex. Analysis of the Rhs2CT-RhsI2 interface using PISA^28^ revealed strong interactions, with an estimated free energy of interface formation (ΔG) of −7.2 kcal/mol, 12 hydrogen bonds, and an interface area of 785 Å^2^ (Figure 2G). These values are indicative of a very stable complex, as expected based on the physiological requirement for immunity proteins to efficiently bind and neutralise their effectors. Consistent with this strong interaction, formation of the Rhs2CT-RhsI2 complex does not require EF-Tu and is readily observed on mixing purified Rhs2CT and RhsI2 proteins (Figure 2H).

### Association of EF-Tu is primarily mediated through an extensive interaction with Rhs2CT and involving the cofactor GDP

The co-purification of EF-Tu with the Rhs2CT-RhsI2 complex was not anticipated and it was important to address the possibility that it might be an artifact. However, structural analysis revealed that the Rhs2CT-EF-Tu interface involves many specific interactions and suggested that the association between Rhs2CT and EF-Tu is likely to be physiologically relevant. Despite the modest estimated free energy change (ΔG of −3.7 kcal/mol) upon Rhs2CT-EF-Tu complex formation, the interface involves a large, buried surface area (890 Å²) and extensive structural interactions, including 17 hydrogen bonds and seven salt bridges, indicating that electrostatic forces contribute to the stability of the complex (Figure 3A). Notably, Rhs2CT interacts with EF-Tu both directly and via the Mg^2+^-GDP cofactor in the active site of EF-Tu. In the absence of GDP, the estimated free energy change of complex formation between Rhs2CT and EF-Tu was reduced by ~0.2 kcal/mol, suggesting that GDP participates in complex formation but has minimal impact on overall binding stability. Importantly, the interaction between Rhs2CT and EF-Tu is sufficient for complex formation in the absence of RhsI2, as shown by the copurification of EF-Tu with catalytically-inactive His_6_-Rhs2CT_CM_ alone (Figure 3B). This indicates that incoming T6SS-delivered Rhs2CT would bind EF-Tu in a recipient target cell.

**Figure 3.**
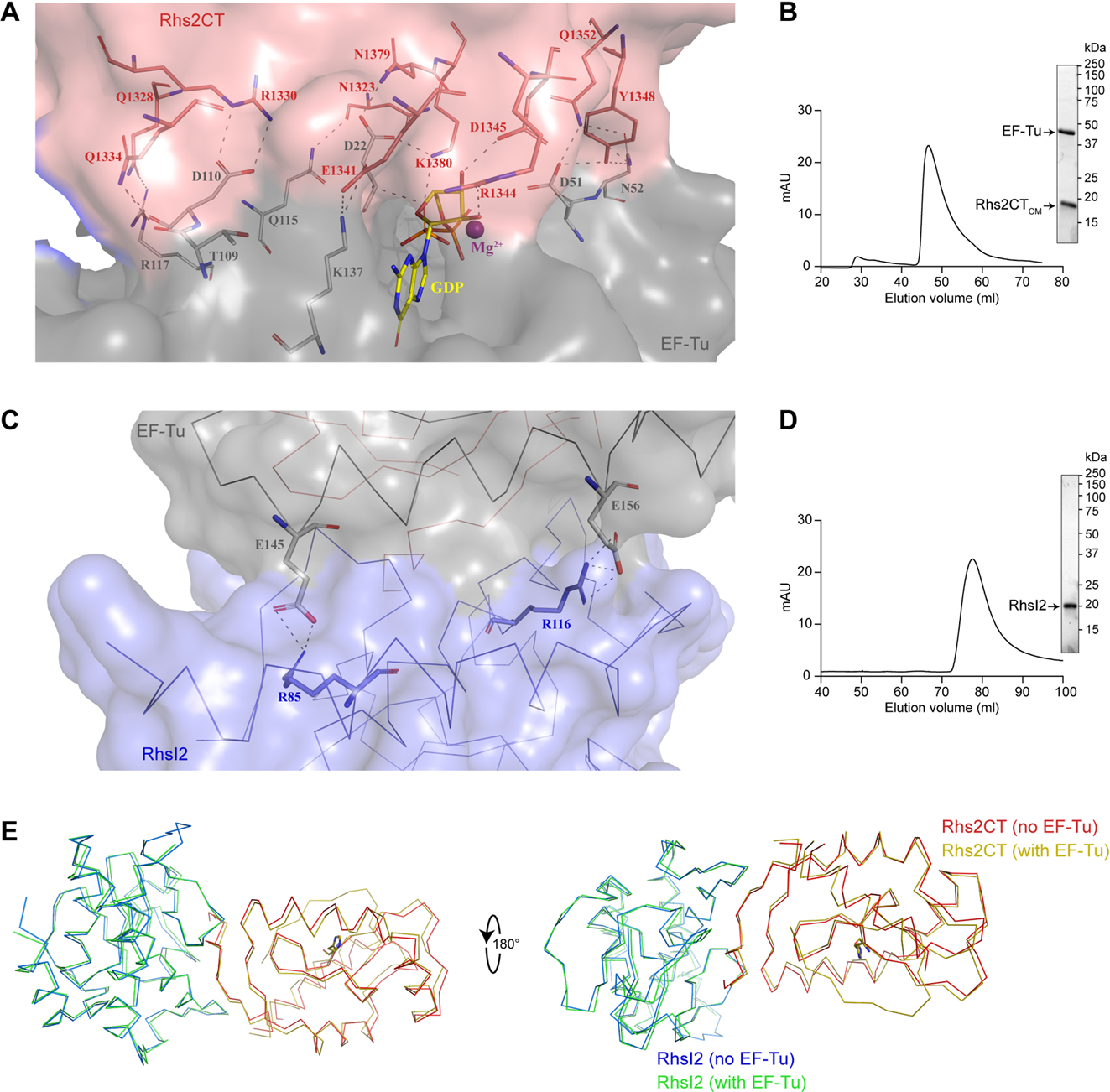
Association of EF-Tu with the Rhs2CT-RhsI2 complex is primarily mediated through an extensive interaction between EF-Tu and Rhs2CT involving the cofactor GDP. **(A)** Close-up depiction of interactions between Rhs2CT (red) and EF-Tu (grey) proteins within the heterotrimeric Rhs2CT-RhsI2-EF-Tu complex. Residues of interest and GDP are shown as sticks, hydrogen bonds (within 2.5 Å-3.5 Å distance) are shown as dashed lines and Mg^2+^ as a purple sphere. For clarity, the Cα backbone is not shown. **(B)** Size-exclusion chromatography, using a Superdex 75 16/60 column, of His_6_-Rhs2CT_CM_ purified from *E. coli*. Proteins from the peak were visualised by SDS-PAGE and Coomassie staining (inset) and the estimated molecular weight corresponding to the peak elution volume is noted. **(C)** Close-up depiction of interactions between RhsI2 and EF-Tu within the Rhs2CT-RhsI2-EF-Tu complex with residues of interest shown as sticks and hydrogen bonds as dashed lines. The Cα backbone is shown for RhsI2 (blue), EF-Tu (grey) and, behind, Rhs2CT (red). **(D)** Size-exclusion chromatography, using a Superdex 75 16/60 column, of His_6_-RhsI2 (D) purified from *E. coli*. **(E)** Overlay of the structure of the Rhs2CT-RhsI2 complex alone (Rhs2CT in red, RhsI2 in blue) or when in complex with EF-Tu (Rhs2CT in mustard, RhsI2 in green); for clarity, EF-Tu is not depicted.

Interestingly, there is also an interaction between RhsI2 and EF-Tu in the heterotrimeric Rhs2CT-RhsI2-EF-Tu complex, with an estimated ΔG of −9.1 kcal/mol suggesting a thermodynamically stable interaction, although the interface (580 Å²) is smaller compared to that between Rhs2CT and EF-Tu (Figure 3C). The presence of five salt bridges indicate that electrostatic forces play a role in stabilising the interaction. However, these alone are unlikely to fully explain such a favourable ΔG, implying that van der Waals interactions and hydrophobic contacts contribute significantly to binding energy. However, this interaction is not sufficient to allow RhsI2 to copurify EF-Tu in the absence of Rhs2CT (Figure 3D). Hence RhsI2 alone does not interact with EF-Tu but it is predicted to enhance the stability of the interaction between Rhs2CT and EF-Tu in the context of the heterotrimeric complex. In order to determine whether interaction with EF-Tu affects the conformation of Rhs2CT or RhsI2, we determined the crystal structure of the Rhs2CT-RhsI2 heterodimeric complex formed from isolated Rhs2CT and RhsI2 proteins to 2.17 Å resolution (Supplementary Table 3). Comparison of the Rhs2CT–RhsI2 complex with the Rhs2CT–RhsI2–EF-Tu heterotrimer revealed that heterotrimer formation does not affect the conformation of Rhs2CT (RMSD 0.61Å) or RhsI2 (RMSD 0.35Å) (Figure 3E, Supplementary Figure 2). Similarly, a comparison of EF-Tu within the heterotrimer with other EF-Tu structures available in the PDB (e.g. PDB 1EFC) demonstrates that the structure of the elongation factor remains largely unaffected by forming a complex with the effector-immunity pair (overall RMSD 1.23Å, with the slightly higher value reflecting inter-domain flexibility of EF-Tu). This suggests that EF-Tu binding does not induce structural rearrangements in the Rhs2CT-RhsI2 complex and likely serves a stabilising role, with the interaction being primarily driven by surface complementarity. The strength of the overall interaction between the Rhs2CT-I complex and EF-Tu is indicated by both thermodynamic (ΔG of −13.2 kcal/mol) and structural parameters, namely a large total interface area (1440 Å²), extensive hydrogen bonding (20 hydrogen bonds), and strong electrostatic forces (12 salt bridges). Taken together, our structural analysis strongly suggests that EF-Tu binding to Rhs2CT and Rhs2CT-RhsI2 is physiologically relevant.

### Interaction of Rhs2CT with EF-Tu is required for T6SS-dependent intoxication of bacterial cells by Rhs2 but is not required for Rhs2CT toxicity per se or for intoxication of fungal cells

Our structural studies revealed a specific interaction between Rhs2CT and EF-Tu, raising the possibility that this interaction may have a functionally-significant role during T6SS delivery or intoxication of recipient cells. In order to investigate this possibility, we needed to prevent the interaction between Rhs2CT and EF-Tu. Since EF-Tu is an essential bacterial protein we could not use mutants lacking EF-Tu, and instead we aimed to generate a mutant of Rhs2CT which could no longer interact with EF-Tu. We used the Rhs2CT-RhsI2-EF-Tu structure to examine the interface between Rhs2CT and EF-Tu and introduce amino acid substitutions into Rhs2CT which should disrupt the interaction between the two proteins. Initial attempts revealed that two or three substitutions were insufficient to prevent the interaction, and a final variant with four substitutions was generated: Rhs2 E1341R, R1344A, A1378W, K1380E (Rhs2CT_EM_, Rhs2 <u>E</u>F-Tu-non-interacting <u>m</u>utant). An additional non-toxic variant containing these substitutions plus the catalytically-inactive mutation (H1369A) was also generated (Rhs2CT_EM-CM_). Pull-down assays using purified Rhs2CT protein immobilised on Ni^2+^-affinity resin showed that wild type Rhs2CT, as expected, efficiently pulled down EF-Tu from total *E. coli* lysate, whilst Rhs2CT_EM-CM_ did not pull down any EF-Tu (Figure 4A), confirming that the four substitutions in Rhs2CT_EM_ abolish a specific interaction with EF-Tu. The Rhs2CT_EM-CM_ protein was used since overproduction of Rhs2CT_EM_ with RhsI2 resulted in a small amount of unbound toxin which was sufficient to cause toxicity to the producing *E. coli* cells and prevent purification of the complex for subsequent Rhs2CT isolation (Supplementary Figure 3). This may imply that the Rhs2CT_EM_-RhsI2 complex is slightly less stable than the native version, but the effect was only seen in this overproduction system where there may be relatively less immunity protein than in more relevant situations. We then determined whether the fungal equivalent of EF-Tu, mitochondrial elongation factor EF-Tu, or any other protein, was pulled down from total *S. cerevisiae* lysate by Rhs2CT. However, no such interaction was detected (Figure 4B). Rhs2CT_CM_ and Rhs2CT_EM-CM_ were also isolated from *E. coli* cells, using the catalytically-inactive version to prevent toxicity. EF-Tu was again co-purified with Rhs2CT_CM_ but not Rhs2CT_EM-CM_, confirming that the interaction of EF-Tu with Rhs2CT is prevented in the Rhs2_EM_ mutant (Figure 4C).

**Figure 4.**
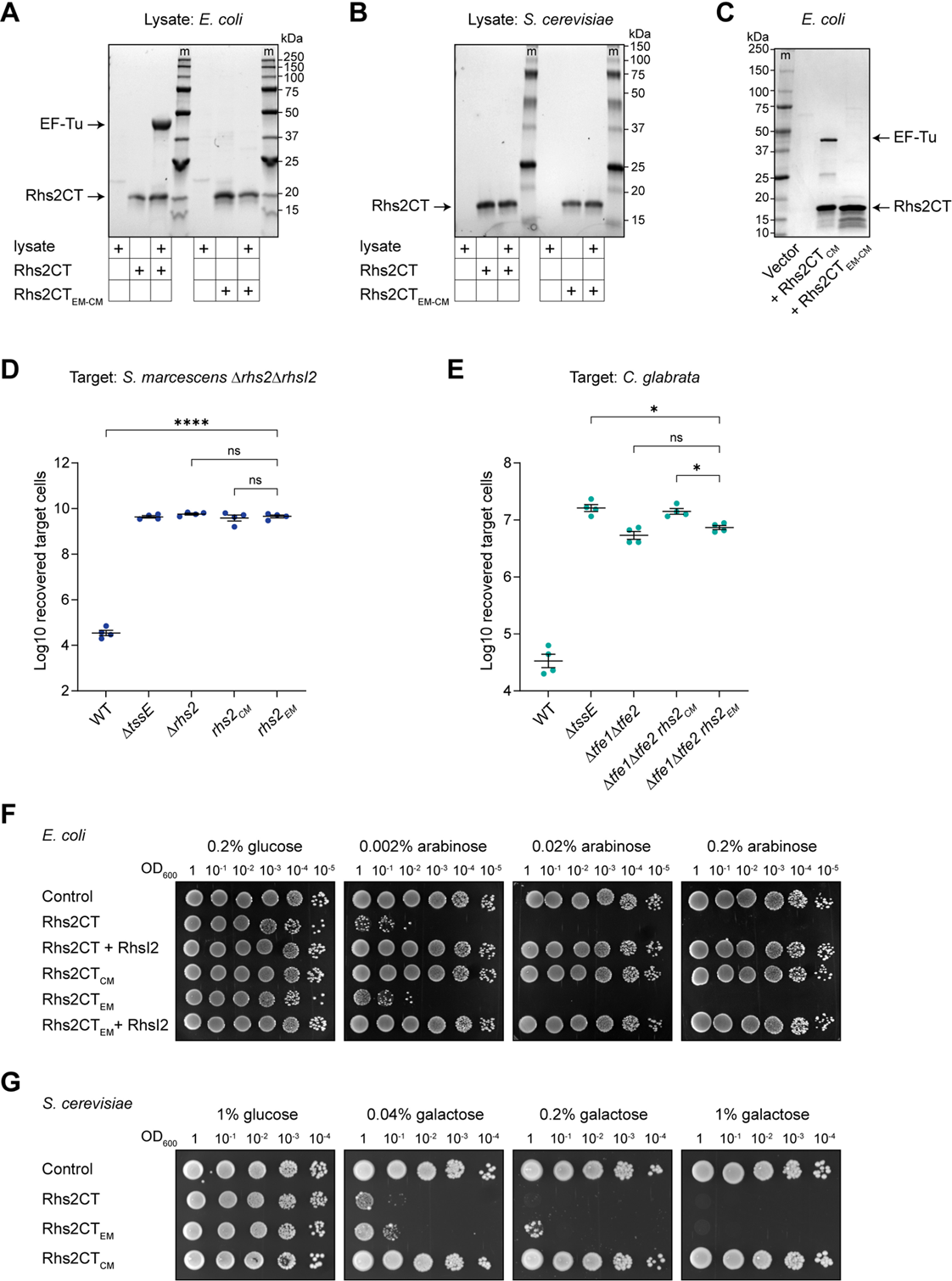
Interaction of Rhs2CT with EF-Tu is required for T6SS-dependent intoxication of bacterial cells by Rhs2 but not for Rhs2CT toxicity per se or intoxication of fungal cells. **(A,B)** Affinity isolation of proteins interacting with Rhs2CT from total lysate samples of *E. coli* MG1655 (A) or *S. cerevisiae* K699 (B). The bait was immobilised wild type His_6_-Rhs2CT (Rhs2CT) or a mutant version (Rhs2CT_EM-CM_) lacking residues involved in EF-Tu interaction (E1341R R1344A A1378W K1380E) and catalysis (H1369A). Control samples were beads with lysate alone or immobilised Rhs2CT protein with no lysate. **(C)** Affinity isolation of Rhs2CT_CM_ (H1369A) and Rhs2CT_EM-CM_ from cells of *E. coli* expressing the His_6_-tagged fusion proteins. In parts A to C, eluate samples were separated by SDS-PAGE followed by Coomassie staining and the positions of protein bands corresponding to Rhs2CT and EF-Tu are indicated (m, molecular weight markers). **(D, E)** Recovery of target bacterial cells (*S. marcescens* Db10 Δ*rhs2*Δ*rhsI2*, part D) or fungal cells (*C. glabrata* ATCC2001, part E), following co-culture with wild type Db10 (WT) or mutants carrying in-frame gene deletions or point mutations in *rhs2* (Rhs2_EM_, Rhs2 E1341R R1344A A1378W K1380E). Data are presented as mean ± SEM with individual data points overlaid (n=4 biological replicates; **** P<0.0001, * P<0.05, ns not significant; one-way ANOVA with Tukey’s test and selected comparisons shown (D) or Šídák’s test (E). **(F)** Growth of *E. coli* MG1655 carrying the empty vector (control) or plasmids directing the expression of Rhs2CT or Rhs2CT_EM_, with or without RhsI2, on media containing glucose or L-arabinose, for repression or induction, respectively, of gene expression. **(G)** Growth of *S. cerevisiae* K699 chromosomal integration strains carrying the empty promoter construct (control), or constructs directing the expression of wild type or mutant versions of Rhs2CT, on media containing glucose or galactose for repression or induction, respectively, of gene expression.

Next, we used the EF-Tu-non-interacting version of Rhs2CT to determine the importance of the Rhs2CT-EF-Tu interaction for Rhs2-mediated, T6SS-dependent antibacterial and antifungal activity. Co-culture assays using the Rhs2-sensitive Δ*rhs2*Δ*rhsI2* target strain revealed that the *S. marcescens* Db10 *rhs2_EM_* mutant showed no activity against susceptible bacterial cells, being indistinguishable from the Δ*rhs2* or *rhs2_CM_* mutants in this assay (Figure 4D). In contrast, antifungal activity was not affected in the *rhs2_EM_* mutant, with the activity of Db10 Δ*tfe1*Δ*tfe2 rhs2_EM_* against *C. glabrata* being the same as Db10 Δ*tfe1*Δ*tfe2* and increased compared with Db10 Δ*tfe1*Δ*tfe2 rhs2_CM_* (Figure 4E). Consistently, the *rhs2_EM_* mutant, exactly like the *rhs2_CM_* mutant, showed no loss of overall T6SS functionality (Figure 1B).

In order to determine whether the loss of Rhs2 antibacterial activity observed in the *rhs2_EM_* mutant upon T6SS-mediated delivery was due to a loss of the toxic enzymatic activity of the CT in bacterial cells, we assessed the toxicity of heterologous expression of variants of Rhs2CT, with and without RhsI2, in *E. coli* cells. Using a tightly-repressed expression system, we observed that expression of Rhs2CT_EM_ is still highly toxic to *E. coli*, almost indistinguishable from wild type Rhs2CT, whilst toxicity of both versions could be completely rescued by co-expression of RhsI2 or in Rhs2CT_CM_ (Figure 4F). Similarly, and consistent with the anti-fungal co-culture assays, Rhs2CT_EM_ retains toxicity when expressed in *S. cerevisiae* (Figure 4G), although a slight decrease in toxicity compared with wild type Rhs2CT may reflect a small loss of stability in the Rhs2CT_EM_ variant.

Taken together, our data show that the interaction between Rhs2CT and EF-Tu is not required for Rhs2 secretion or for the stability or toxic enzymatic function of Rhs2CT, in either bacterial or fungal cells. However, it is essential for successful T6SS-dependent intoxication of bacterial, but not fungal, cells. This implies that the Rhs2CT-EF-Tu interaction plays a role in an aspect of the delivery process unique to bacterial target cells, such as the requirement to cross a second membrane in order to reach the cytoplasm.

### Rhs2CT degrades DNA and localises to the nucleus in fungal cells

Next, we aimed to investigate the basis of Rhs2CT toxicity in fungal cells. In this case, in contrast with bacterial cells, DNA is confined within membrane-enclosed organelles rather than accessible in the cytoplasm and EF-Tu is not available to assist trafficking. Initially, we confirmed that Rhs2CT exerts DNase activity and degrades DNA in fungal cells. Expression of an EGFP-Rhs2CT fusion in *S. cerevisiae*, but not EGFP alone, caused DNA breaks detectable using a TUNEL assay (Figure 5A,B). Fusion of EGFP to the C-terminus of Rhs2CT did not impair toxicity (Supplementary Figure 4). This result implied that Rhs2CT must be able to access DNA in the nucleus or the mitochondria of the fungal cells. Therefore, we used fluorescence microscopy to determine the localisation of EGFP-Rhs2CT_CM_ expressed in *S. cerevisiae*, using the catalytically inactive mutant to prevent toxicity and ensure comparable growth and cellular integrity with the EGFP-only control. The EGFP-Rhs2CT_CM_ fusion protein was confirmed to be intact and stable by immunoblotting (Supplementary Figure 4). Visualisation of *S. cerevisiae* expressing EGFP alone showed EGFP signal distributed throughout the cells apart from the vacuole. In contrast, EGFP-Rhs2CT_CM_ signal was concentrated in the nucleus, as shown by colocalisation with the nuclear stain NucBlue, whilst no association of EGFP-Rhs2CT_CM_ with mitochondria (stained with MitoFix640) was observed (Figure 5C, Supplementary Figure 5). Quantitative colocalisation analysis confirmed strong colocalisation of EGFP-Rhs2CT_CM_, but not EGFP, with NucBlue (Figure 5D). EGFP-Rhs2CT_CM_ showed even lower colocalisation with mitochondria than EGFP alone, reflecting its movement from the cytoplasm to the nucleus. Therefore, Rhs2CT does indeed act as a DNase toxin in fungal cells and is localised to the nucleus, implying it acts on nuclear DNA. These data also suggest that Rhs2CT has the intrinsic capacity to translocate into the fungal nucleus from the cytoplasm, rather than requiring the T6SS to deliver it into the nucleus. This agrees with the observation that expression of RhsI2 in the cytoplasm of yeast cells can provide protection from either T6SS-delivered or heterologously-expressed Rhs2CT (Figure 1). The sensor kinase Mec1 is a central component of the DNA damage checkpoint which detects nuclear DNA damage and activates the DNA damage response. Mutants of *S. cerevisiae* lacking *MEC1* are non-viable but can be rescued by deletion of *SML1*, with the double mutant being viable but more sensitive to DNA damage^29, 30^. Consistent with Rhs2CT being a T6SS-delivered DNase which acts on nuclear DNA, a *mec1*Δ*sml1*Δ mutant of *S. cerevisiae* was more susceptible to Rhs2CT than the parental strain, showing reduced recovery when co-cultured with strains of *S. marcescens* able to deliver wild type Rhs2CT (Fig 5E).

**Figure 5.**
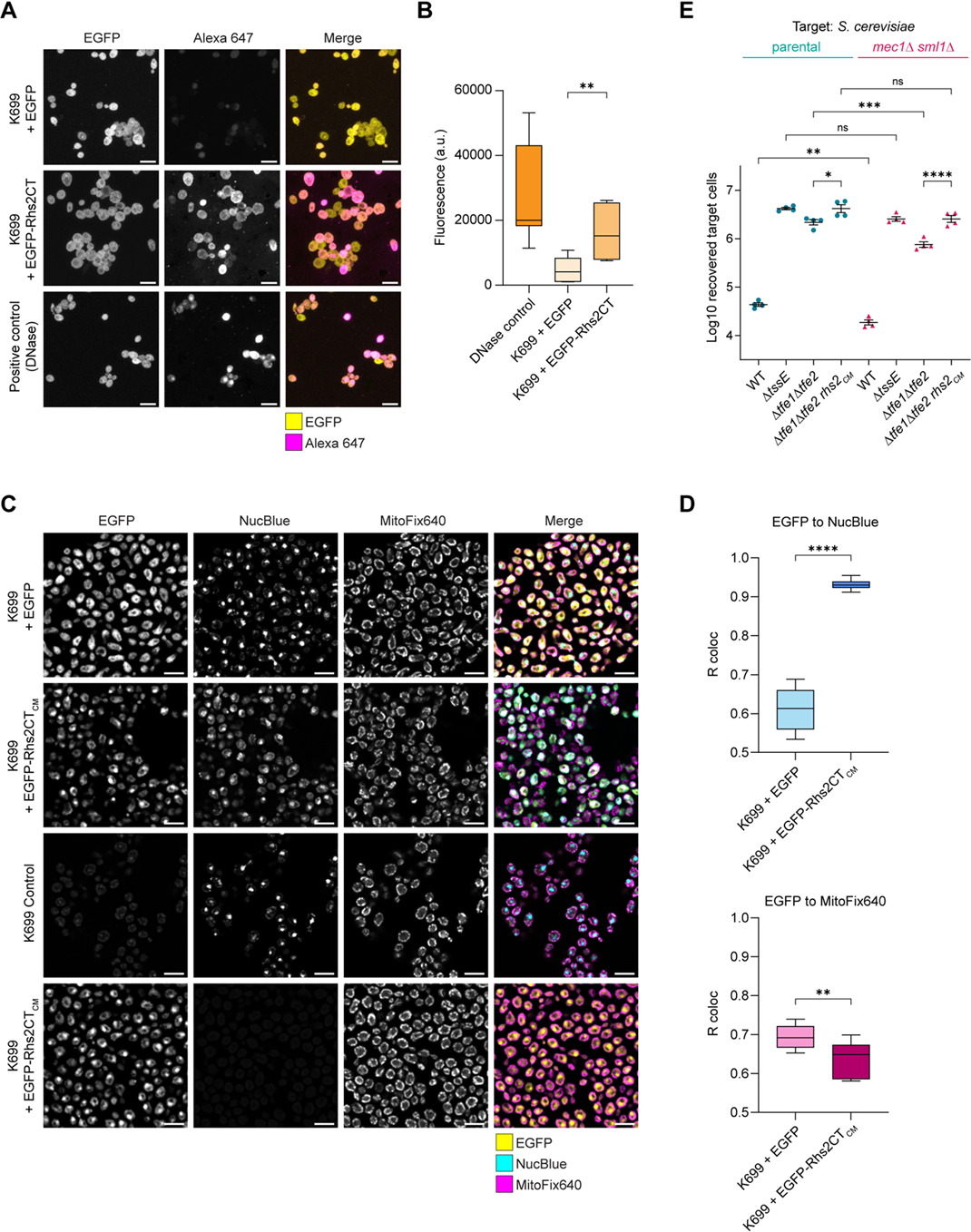
Rhs2CT generates breaks in DNA and localises to the nucleus in fungal cells. **(A)** Representative images from TUNEL assay performed on cells of *S. cerevisiae* K699 expressing EGFP alone (EGFP) or EGFP fused to the N-terminus of wild type Rhs2CT (EGFP-Rhs2CT) from chromosomally-integrated expression constructs, shown as single fluorescent channels of EGFP and TUNEL (Alexa 647) and the merged channels (Merge). **(B)** Quantitative comparison of relative fluorescence intensity values from the cellular TUNEL assays. Data are presented as box plots ± SEM (10-15 fields of view for each condition from n=3 biological replicates); **** P<0.0001; one-way ANOVA with Šídák’s test. **(C)** Representative single slice images of K699 expressing EGFP alone or EGFP fused to the N-terminus of catalytically-inactive Rhs2CT (EGFP-Rhs2CT_CM_, H1369A), shown as single channels of EGFP, NucBlue and MitoFix 640 and merged channels. In the bottom row, NucBlue staining was not performed to allow clearer comparison between EGFP and MitoFix 640. **(D)** Quantification of fluorescence colocalisation values between the indicated channels was performed using Pearsons correlation with Costes thresholding. Data are presented as box plots ± SEM (7-13 fields of view for each condition from n=3 biological replicates); **** P<0.0001, ** P<0.01; one-way ANOVA with Šídák’s test. All scale bars are 5 μm. **(E)** Recovery of *S. cerevisiae* W303-1a (parental) and its derivative lacking *MEC1* and *SML1* (*mec1*Δ *sml1*Δ) following co-culture with wild type *S. marcescens* Db10 or mutants carrying in-frame gene deletions or point mutation in *rhs2* as indicated. Data are presented as mean ± SEM with individual data points overlaid (n=4 biological replicates; **** P<0.0001, *** P<0.001, ** P<0.01, * P<0.05, ns not significant; one-way ANOVA with Tukey’s test; for clarity, only selected comparisons are displayed).

### Rhs2CT uses the nuclear import machinery to access the fungal nucleus

The previous data indicated that Rhs2CT can access the nucleus of fungal cells by translocation from the cytoplasm, rather than requiring direct delivery by the T6SS. Therefore we predicted that Rhs2CT uses the nuclear import machinery, despite not displaying a classical nuclear localisation signal (NLS). To test this hypothesis, we used a temperature-sensitive nuclear import mutant of *S. cerevisiae* (importin-α *srp1-31*, strain NOY612). A conditional mutant was required due to the essentiality of the nuclear import machinery and the *srp1-31* allele was selected since it displays a nuclear import defect but not a proteasome targeting defect^31^. NOY612 was transformed with the same chromosomal integration constructs directing galactose-inducible expression of Rhs2CT, EGFP and EGFP-Rhs2CT_CM_ as used above in K699, in order to examine toxicity and localisation of Rhs2CT when the function of the nuclear import machinery was compromised. At 23°C, we observed a slight growth impairment in NOY612 expressing EGFP or non-toxic Rhs2CT_CM_ compared with K699, implying some reduction in importin-α function. In contrast, NOY612 expressing toxic Rhs2CT survived better than K699 expressing Rhs2CT at low induction levels (0.04% galactose, Figure 6A). At 30°C, when importin-α function is more severely compromised, NOY612 strains expressing EGFP or Rhs2CT_CM_ were more severely impacted for growth compared with K699 (but not completely unable to grow as they would be at 37°C). However, strikingly, despite displaying reduced growth overall, NOY612 cells were more resistant to Rhs2CT than K699, indicating that Rhs2CT was indeed less able to access the nucleus and cause DNA damage when nuclear importin function was reduced (Figure 6A).

**Figure 6.**
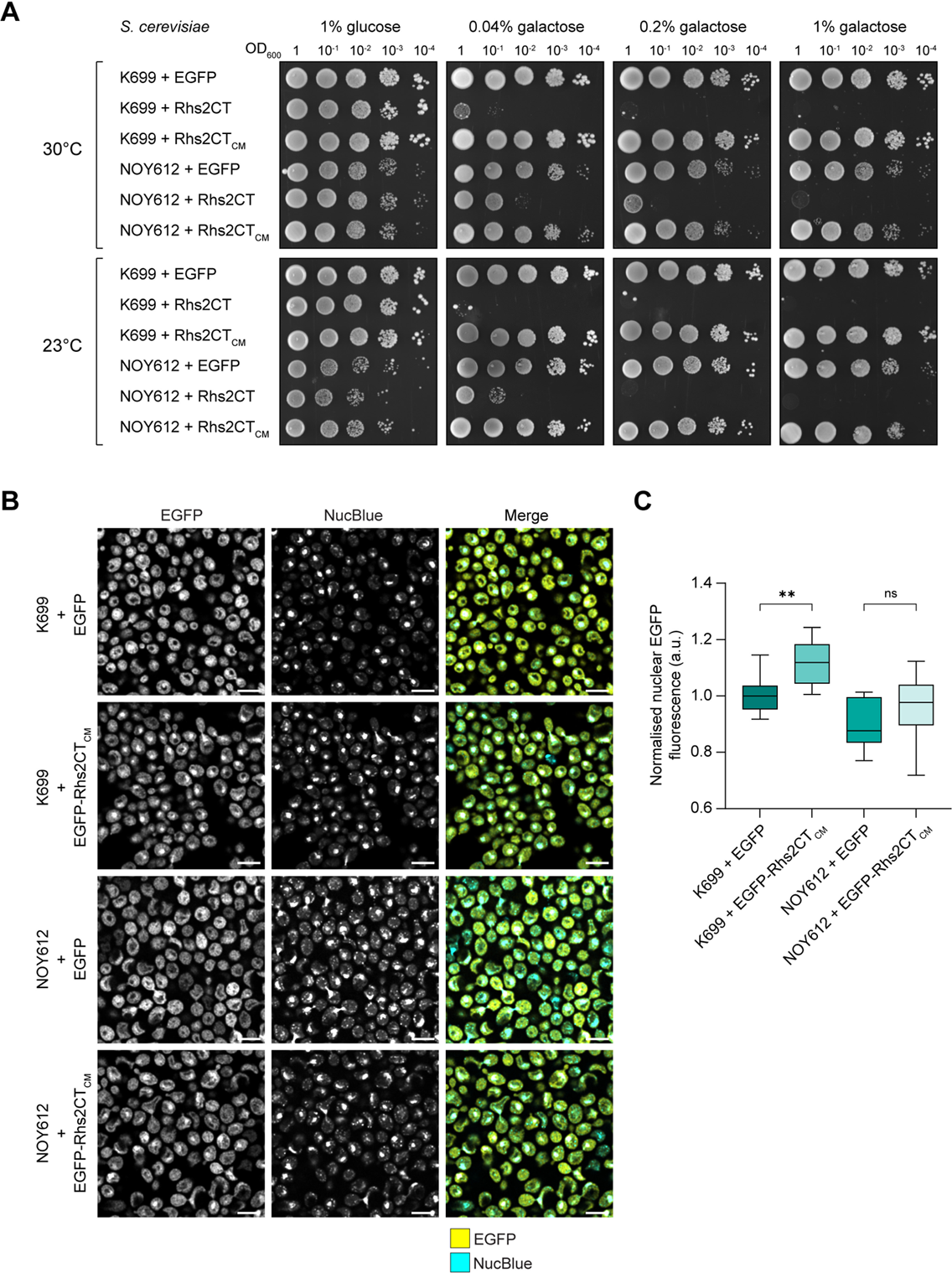
The ability of Rhs2CT to access the nucleus of fungal cells is impaired in temperature-sensitive nuclear import mutants. **(A)** Growth of *S. cerevisiae* K699 or NOY612 (importin-α mutant *srp1-31*) carrying chromosomally-integrated constructs directing the expression of EGFP (control), the wild type Rhs2 C-terminal domain (Rhs2CT) or a catalytically inactive variant (EGFP-Rhs2CT_CM_, H1369A) on media containing glucose or galactose for repression or induction, respectively, of gene expression. Growth was assessed at 30°C (top panel) and 23°C (bottom panel). **(B)** Representative single slice images of *S. cerevisiae* K699 or NOY612 carrying constructs directing the expression of EGFP alone (EGFP) or EGFP fused with Rhs2CT_CM_ (EGFP-Rhs2CT_CM_) shown as single channels of EGFP and NucBlue and merged channels (Merge). Expression was induced using 1% galactose at 30°C. (**C)** Quantification of EGFP nuclear fluorescence intensity normalised to cytoplasmic EGFP signal. Data are presented as box plots ± SEM (14-17 fields of view for each condition from n=3 biological replicates); ** P<0.01, ns not significant; one-way ANOVA with Šídák’s test. All scale bars are 5 μm.

To determine directly whether nuclear localisation was reduced in the temperature-sensitive nuclear import mutant, EGFP and EGFP-Rhs2CT_CM_ were visualised in K699 and NOY612 during growth at 30°C. This revealed that some of the NOY612 cells had irregular shapes and that nuclear localisation of EGFP-Rhs2CT_CM_ was discernible in some cells but was overall less obvious (Figure 6B). The amount of EGFP fluorescence colocalising with NucBlue (localised to the nucleus) relative to the EGFP fluorescent signal in the rest of the cell was determined in each case. This showed that there was significantly higher nuclear fluorescence for EGFP-Rhs2CT_CM_ than EGFP in K699, but not in NOY612 (Figure 6C). Taken together, these data show that compromising nuclear import leads to reduced toxicity and nuclear localisation of Rhs2CT, consistent with this import machinery being used for translocation of Rhs2CT and EGFP-Rhs2CT into the fungal nucleus.

## DISCUSSION

Here, we have shown that Rhs2 of *S. marcescens* Db10 is a dual-kingdom effector whose C-terminal DNase domain hijacks distinct, essential target cell functions in order to reach its site of action in bacterial and fungal cells. In bacterial target cells, incoming T6SS-delivered Rhs2CT interacts with EF-Tu to allow it to access DNA within the cytoplasm. In the case of fungal target cells, where Rhs2 represents the remaining antifungal effector of *S. marcescens* Db10, Rhs2CT uses the nuclear import machinery to access DNA sequestered in the membrane-bound nucleus. These findings reveal that whilst T6SS effectors may frequently have the enzymatic capacity to act as dual-kingdom effectors, specific adaptations may be required to allow them to exert toxicity against both types of target cell.

Our structural and *in vitro* analyses confirmed that the Rhs2CT is a DNase toxin and revealed that it is an atypical member of the HNH endonuclease superfamily. Rhs2 can be a very potent effector upon T6SS-mediated delivery into bacterial competitors, causing a dramatic reduction in recovery of susceptible target cells despite being delivered at a rate of less than one molecule per firing event of the T6SS (since it represents only one of three alternative PAAR tip proteins^16^). DNase enzymes can be very effective toxins since they will be lethal if an intoxicated cell cannot repair the multiple breaks generated in its genomic DNA. This is likely the reason why DNases are common secreted anti-bacterial toxins across multiple systems including Cdi, T4SS and T6SS^5, 32, 33^. Unusually for T6SS effector-immunity pairs^5^, the cognate immunity protein, RhsI2, does not neutralise the Rhs2CT toxin by blocking its active site, but instead binds elsewhere on the protein. It is possible to speculate that this mode of interaction provides more evolutionary freedom for the CT and immunity protein to co-evolve, for example to become effective against competitor strains carrying the original immunity protein, without compromising the enzymatic function of the toxin. More broadly, acquisition and evolution of new T6SS effector and immunity proteins is believed to lead to ‘arms races’ between different bacterial lineages as they compete with each other^1, 3^.

This study has revealed a further aspect of how Rhs2 has evolved to function as an antibacterial effector, namely its requirement to interact with the essential protein EF-Tu in bacterial target cells. Our data suggest that this interaction is required for the incoming CT to access the cytoplasm of the target cells. In general, the mechanism by which RhsCT domains delivered by the T6SS are released from the Rhs β-barrel ‘shell’ and cross the inner membrane of intoxicated bacterial cells remains to be defined. Indeed, it may differ depending on the N-terminal domain of the Rhs protein, with a number of different N-terminal domains which include, or mediate interaction with, PAAR or VgrG domains having now been identified^34, 35^. In all cases, the RhsCT is cleaved by autoproteolytic cleavage within the shell and initially remains trapped inside in an unfolded or molten globule conformation^12, 13, 34, 35, 36, 37^. However, where and how the shell opens to release the CT is unclear and may vary. In the most common class of T6SS-associated Rhs proteins, which includes Rhs2, the N-terminal domain contains a prePAAR motif and transmembrane helix-containing domain (TMD) upstream of a PAAR domain^11,35^. This TMD, which is stabilised by Eag chaperones prior to secretion, has been implicated in allowing the incoming RhsCT to cross the inner membrane^11^, although several distinct models have been proposed for how this may occur. In one model, the incoming T6SS spike reaches the periplasm and the CT is ‘tugged’ out of the shell and threaded across the IM at the site of insertion of the TMD. Subsequent folding of the CT in the cytoplasm drives the translocation across the inner membrane^12^. Alternatively, it has been proposed that the TMD may act as a ‘molecular grease’ to facilitate translocation of the incoming T6SS spike, including the Rhs shell, right across the inner membrane to the cytoplasm^11^, or that the shell and CT may be translocated across the inner membrane with the assistance of the TMD region^12, 13^. In these cases, release of the CT from the shell in the cytoplasm is again likely to be driven by its folding. We propose that interaction of Rhs2CT with EF-Tu, together with folding of the CT after its release from the shell, provides the requisite driving force for the Rhs2CT to be imported across the bacterial inner membrane, with or without the shell. We believe that our data showing that this interaction is not required for translocation into fungal cells is consistent with a model involving a specific import mechanism across the bacterial inner membrane.

Rhs2 represents the first example of an Rhs C-terminal domain requiring a target cell protein for successful intoxication due to a role in translocation of the CT. Another RhsCT, the ADP-ribosyltransferase TreX from *Xenorhabdus*, was also shown to require interaction with a target cell protein, thioredoxin 1, to exert toxicity^38^. However, in contrast, this interaction was required for the stability and activity of the toxin, although it may also drive translocation to the cytoplasm in an analogous manner to the EF-Tu-Rhs2CT interaction. Interestingly, a very similar mechanism for EF-Tu-driven import was proposed for the unrelated T6SS NADase effector Tse6 from *Pseudomonas aeruginosa*, where interaction of Tse6 with EF-Tu was required for Tse6 to be delivered into target cells^39^. Whilst the authors were unable to determine whether Tse6 interacts with EF-Tu in attacker cells, target cells, or both, in our system we can conclude that Rhs2CT interacts with EF-Tu exclusively in target cells since the CT is sequestered within the Rhs shell prior to secretion. EF-Tu is a GTPase which delivers charged aminoacyl-tRNAs to the ribosome during translation elongation^40^. The fact that it is an essential, highly conserved and abundant housekeeping protein makes it both a good target and a good hijacked helper protein for antibacterial toxins. For example, in addition to allowing import of T6SS-delivered Rhs2CT and Tse6, interaction with EF-Tu is also required for the enzymatic activity of two CdiA C-terminal toxin domains^41, 42^, whilst the RhsCT, Tre^Tu^, and the T-AT toxin, Doc, inhibit protein synthesis by ADP-ribosylating or phosphorylating, respectively, EF-Tu^43, 44^. It will be interesting to discover how many other RhsCTs also need to bind target cell proteins to drive their import across the inner membrane and what properties of the T6SS, Rhs core and/or CT shape this requirement.

This study has also revealed a second role for Rhs2, as an antifungal effector delivered by the T6SS, showing that Rhs effectors can be deployed against fungal cells. As a large specialised effector, with dual antibacterial and antifungal function, Rhs2 presents a striking contrast with the other antifungal T6SS effectors delivered by *S. marcescens* Db10, Tfe1 and Tfe2, which are small Hcp-dependent cargo effectors with activity specific to fungal cells^7, 16^. It was somewhat unexpected for the missing antifungal effector of Db10 to be an antibacterial DNase, given the need for such an activity to access the interior of a membrane-bound organelle rather than, for example, degrade NAD, RNA or phospholipids which are more accessible. However, we have demonstrated that Rhs2CT does indeed have the intrinsic ability to access the nucleus and appears to make use of the nuclear import machinery. The nuclear envelope forms an impermeable barrier and proteins must enter the nucleus via numerous nuclear pore complexes. Movement through these complexes occurs via facilitated diffusion mediated by nuclear transport factors which recognise nuclear transport signals, or, alternatively, molecules without transport signals can enter the nucleus by passive diffusion, which is limited by increasing size up to ~40-60 kDa^45^. Rhs2CT is small enough, ~16 kDa, to be able to enter the nucleus by diffusion. However, several lines of evidence suggest that this is not the primary mechanism. Firstly, an EGFP-Rhs2CT fusion (43 kDa) remains toxic and becomes localised to the nucleus far more efficiently than EGFP alone (27 kDa) (Figure 5). Secondly, our data suggest that compromising Srp1-mediated nuclear import decreases toxicity and nuclear localisation of Rhs2CT, implying that nuclear transport machinery does promote Rhs2CT localisation to its site of action. Srp1 is a member of the importin-α family of nuclear transporters that bind proteins with an NLS and is the sole importin-α protein in *S. cerevisiae*^31^. Rhs2CT does not contain a detectable classical NLS, suggesting that it might interact with a yeast protein which does have an NLS in order to ‘piggy back’ into the nucleus^46^. The relatively modest impact of Rhs2 in competition with fungal cells, compared with its potency against bacterial competitors, could be due to the need for the CT to further translocate from the cytoplasm into the nucleus, but may also be due to fewer T6SS firing events being ‘on target’ (penetrating the thick fungal cell wall) and/or partial protection of the DNA by packaging into chromatin.

The antifungal activity of Rhs2 was discovered following the observation that a Δ*tfe1*Δ*tfe2* mutant of *S. marcescens* Db10 retains activity against *C. glabrata* and *S. cerevisiae* compared with the T6SS-inactive Δ*tssE* mutant. Interestingly, however, the Δ*tfe1*Δ*tfe2* mutant shows no T6SS-dependent antifungal activity against *Candida albicans^7^*, implying that *C. albicans* is resistant to Rhs2. The reason for this difference is not yet clear but several hypotheses can be proposed: first, Rhs2CT, for unknown reasons, cannot be released from the Rhs shell or cannot utilise the nuclear import machinery in *C. albicans*; second, *C. albicans* can recover more efficiently from breaks in its genomic DNA caused by Rhs2CT than the other species. We believe that *C. albicans* is resistant to Rhs2 because it is better able to survive DNA damage compared with *C. glabrata* and *S. cerevisiae*. In support of this idea, a *mec1*Δ mutant of *C. albicans*, which is impaired in the ability to respond to nuclear DNA damage^47^, is sensitive to T6SS-delivered Rhs2 (Supplementary Figure 6). *Candida glabrata* is an opportunistic pathogen more closely related to *S. cerevisiae* than *C. albicans.* Clinical isolates display high variability in chromosome structure and rapidly evolve antifungal drug resistance^48^. It has been reported that *C. glabrata* cells exposed to DNA damaging agents such as the alkylating agent methyl methanesulfonate (MMS) proceed to S phase and cell division, resulting in loss of viability due to aberrant mitosis^48^. This non-canonical fungal DNA damage response could contribute to its rapid genetic change and drug resistance but also leave it vulnerable to Rhs2. In contrast, the *C. albicans* DNA damage response is very efficient and critical for its pathogenicity^49^, with genotoxic stress agents activating the DNA damage checkpoint and inducing filamentous growth. The base excision repair pathway of *C. albicans* is more efficient than that of *S. cerevisiae* and *C. albicans* can tolerate loss of the DNA damage checkpoint kinase Rad53 whereas *S. cerevisiae* cannot, exemplifying differences in DNA damage response between species^50, 51^. Overall, these findings indicate that fungal cells may be able to use stress and damage responses to provide resistance towards T6SS effectors, a protection mechanism also seen in bacterial cells^52^.

This study has revealed that Rhs2 from *S. marcescens* Db10 is a dual-kingdom (or dual-function) antibacterial and antifungal effector. Other examples of dual-kingdom effectors have been reported, including several phospholipases and a DNase which act against bacterial and mammalian cells, and an effector which induces mitochondrial fragmentation in yeast and mammalian cells^5, 53, 54^. Another antifungal effector which has DNase activity has recently been described, but no evidence of T6SS-dependent activity against bacterial competitors was reported, so it remains to be determined whether this could also be a dual-kingdom effector, and its mechanism of nuclear import was not investigated^55^. Our detailed analysis has demonstrated how dual anti-bacterial and anti-fungal activity can be achieved in a T6SS-delivered effector, by co-opting different target cell functions to assist in each case. This highlights both versatility and economy of the T6SS and its effectors, and is particularly impressive given that the Rhs2CT comprises only 141 amino acids. Our findings, coupled with the other reports of dual-kingdom effectors, imply that dual-kingdom antibacterial-antifungal T6SS effectors may be more widespread than currently appreciated. However, not all antibacterial effectors have a dual function, given that Rhs1 and six antibacterial cargo effectors^16^ do not contribute to the *S. marcescens* Db10 antifungal T6SS activity as far as we can tell. In some cases it is obvious why not (e.g. the peptidoglycan amidases Ssp1 and Ssp2 target the bacterial cell wall^24^) in others, for example Rhs1 whose CT has NADase activity^22^, fungal cells may lack a specific bacterial protein or cellular environment required for delivery or activity.

In conclusion, this study has revealed that Rhs effectors can be delivered into bacterial and fungal cells by the T6SS and that the CT domain can reach its site of action by hijacking functions specific to each type of target cell in order to allow intoxication of competitors from both kingdoms. Given the wide occurrence of T6SS-associated Rhs proteins, they may represent a common strategy by which bacteria compete against fungal cells in polymicrobial communities. Whilst the T6SS is very versatile in its ability to deliver effectors, factors in the target cell can be essential for successful intoxication.

## MATERIALS AND METHODS

### Bacterial strains, plasmids and culture conditions

Bacterial strains and plasmids used in this study are detailed in Supplementary Table 1. Mutant strains of *S. marcescens* Db10 were generated by an allelic exchange using the suicide vector pKNG101^20^. Derivatives of the pRSFDuet-1 and pET15b-TEV plasmids were generated for protein overproduction and purification, whilst plasmids for arabinose-inducible gene expression in bacterial cells, including for assessment of toxicity, were derived from pBAD18-Kn. Details of oligonucleotide primers or synthetic DNA fragments used in plasmid construction are provided in Supplementary Table 2. Unless stated otherwise, bacterial cultures were grown in LB (10 g L^−1^ tryptone, 5 g/L yeast extract, 10 g/L NaCl, with 18 g/L Select agar for solid media), at 30°C for *S. marcescens* and 37°C for *E. coli*. When required, media were supplemented with antibiotics: carbenicillin (Ap) 100 µg ml^−1^, kanamycin (Kn) 50 µg ml^−1^, and streptomycin (Sm) 100 µg ml^−1^; to maintain repression of proteins expressed from pBAD18-Kn, 0.5% glucose was added to the media for cloning and maintenance.

### Fungal strains, plasmids and culture conditions

Fungal strains and plasmids are used in this study are detailed in Supplementary Table 1. Strains of *S. cerevisiae* K699 carrying chromosomal insertions of genes under the control of the P*_GAL1_* promoter were generated using linearised plasmids derived from vector pSC1384 and integrated by allelic exchange into the *HIS3* locus. To delete *MEC1* in *C. albicans*, disruption cassettes containing *ARG4* or *HIS1* nutritional marker genes, flanked by *loxP* sites and 100 bp 5′ and 3′ of the *MEC1* open reading frame, were generated by PCR using the plasmid template pLAL or pLHL^56^. SN148 wild-type cells^57^ were sequentially transformed with deletion cassettes to disrupt both alleles of *MEC1.* Uridine prototrophy was restored by integrating CIp10 at the *RSP1* locus, generating the *mec1Δ* (JC1549) strain. Fungal strains were cultured at 30°C in YPDA (10 g/L yeast extract, 20 g/L peptone, 40 mg/L adenine hemisulfate) or synthetic complete media (SC; 6.9 g/L yeast nitrogen base complete with amino acids, with adenine at 100 mg/L) containing 2% glucose, with 20 g/L Select agar for solid media. If utilising auxotrophies, SC media lacking the respective amino acid or nucleotide was used. For induction of the P*_GAL1_* promoter, *S. cerevisiae* cultures were pre-incubated in SC media lacking leucine (SC-LEU) and containing 2% raffinose, followed by induction of gene expression by the inclusion of 1% galactose, unless stated otherwise.

### Recombinant protein production and purification for crystallography

For production and purification of the Rhs2CT-RhsI2-EF-Tu complex, genes encoding Rhs2CT and RhsI2 were cloned into the two multiple cloning sites of pRSFDuet-1 to generate pSC3502, encoding Rhs2CT with an N-terminal hexa-histidine (His_6_) tag cleavable by Tobacco Etch Virus (TEV) protease and untagged RhsI2. The buffers employed for purification were buffer *A1* (50 mM MES-HCl 50 mM imidazole-HCl pH 5.5, 250 mM NaCl, 1 mM MgCl_2_ and 0.5 mM TCEP) and buffer *B1* (buffer *A* with 0.5 M imidazole). Cells of *E. coli* BL21(DE3) freshly transformed with pSC3502 were used to inoculate 20 ml of LB + Kn and grown for 3 h at 37 °C. This culture was used to inoculate 1 L of Auto-Induction Terrific Broth with Trace Elements and 2% glycerol (AIM TB TE 2% glycerol) media, followed by growth for ~48 h at 22°C. Cells were harvested by centrifugation (4200*g* for 30 min at 4°C). The cell pellet was resuspended in buffer *A* supplemented with complete EDTA-free protease inhibitor cocktail (Thermo Scientific) and DNase I (Sigma Aldrich), then passed through a Cell Disrupter (Constant Systems) at 207 MPa and the lysate clarified by centrifugation at 40 000*g* for 30 min at 4°C. The resulting supernatant was filtered (0.2 µm) and loaded onto a 5 ml HisTrap HP column (Cytiva) previously charged with Ni^2+^ and pre-equilibrated with buffer *A* for the initial affinity chromatography step. The column was washed to remove all unbound proteins, and a linear gradient of buffer *B* was then applied. The protein complex eluted at approximately 300 mM imidazole. Fractions containing the complex were identified using SDS-PAGE, pooled and treated with His-tagged TEV protease (1 mg of protease per 10 mg of protein) overnight at 4°C whilst also being dialysed against buffer *A* to remove excess imidazole. This was followed by reverse affinity chromatography on the HisTrap HP column to remove the TEV protease, cleaved tag and any uncleaved protein that remained. A final purification step was achieved by size-exclusion chromatography (SEC) using a calibrated Superdex 75 HiLoad 26/60 column (Cytiva) in buffer *C1* (25 mM MES-HCl 25 mM imidazole-HCl pH 5.5, 150 mM NaCl, 1 mM MgCl_2_ and 0.5 mM TCEP). Calibration was performed using molecular weight standards: blue dextran (> 2,000 kDa), thyroglobulin (669 kDa), ferritin (440 kDa), aldolase (158 kDa), conalbumin (75 kDa), ovalbumin (43 kDa), carbonic anhydrase (29.5 kDa), ribonuclease A (13.7 kDa) and aprotinin (6.5 kDa) (GE Healthcare). The theoretical mass of the Rhs2CT-RhsI2-EF-Tu*_Ec_* complex is approximately 79.2 kDa. The SEC profile was well defined and suggested the presence of a single species with approximate molecular mass 52 kDa. Peak fractions were pooled and concentrated using centrifugal force (Sartorius spin concentrator). The high level of sample purity was confirmed by SDS-PAGE using stain-free gels (Bio-Rad) prior to crystallisation trials. The yield of Rhs2CT-RhsI2-EF-Tu was ~ 30 mg/L^−1^. Protein concentrations were determined from the absorbance at 280 nm using the Beer– Lambert law with the predicted molar extinction coefficient (ɛ = 72,310 M ^−1^ cm^−1^ at 280 nm) obtained from ProtParam^58^.

Unless specified otherwise, other protein purifications were performed following the method above. Details of each purification, including expression plasmids, buffers, extinction coefficients, and purification steps, is provided in Supplementary Table 4. Lysis/wash buffers are named *A*, elution buffers *B* and SEC buffers *C*, with variants (e.g. *A1*, *B1* etc.) specific to each purification.

### Isolation of Rhs2CT by refolding

Following the initial affinity purification step as above, concentrated His_6_-Rhs2CT-RhsI2-EF-Tu complex was dialysed against buffer *A1* and dissolved in 6 M guanidine-HCl. The mixture was incubated at 90 °C for 5 min with stirring and then manually loaded on a Ni^2+^-charged HisTrap HP column that had been pre-equilibrated with 6 M guanidine-HCl. The column was then sequentially syringe-washed with 6M guanidine-HCl (to remove RhsI2 and EF-Tu), buffer *A2* (to refold the Rhs2CT) and buffer *B2* (to elute refolded Rhs2CT). Refolded Rhs2CT was then subjected to SEC using a pre-equilibrated Superdex 75 HiLoad 16/60 column (Cytiva) pre-equilibrated in buffer *C2* to remove trace amounts of RhsI2 and EF-Tu.

### Reconstitution of Rhs2CT-RhsI2 complex

To obtain the Rhs2CT-RhsI2 complex without EF-Tu, His_6_-RhsI2 was purified alone following the protocol provided above and then combined at near-equimolar quantity with refolded Rhs2CT. This was followed by SEC to remove excess RhsI2 and confirm the oligomeric state of the complex.

### AlphaFold modelling

Models of Rhs2CT and RhsI2 were generated using AlphaFold2^26^ via ColabFold hosted at: https://colab.research.google.com/github/sokrypton/ColabFold/blob/main/AlphaFold2.ipynb. Models were generated using AlphaFold2 (versions v2.0 to v2.3), with multiple sequence alignments (MSAs) produced by the MMseqs2-based pipeline implemented in ColabFold. For each target, five models were generated using default settings. The model with the highest overall pLDDT confidence score was selected for downstream analysis and interpretation.

### Crystallisation, X-ray data collection, processing and refinement

#### Rhs2CT-RhsI2-EF-Tu complex

An initial crystallisation condition was identified in the Nextal screen (Hampton research; HR2-134) following a two week incubation at 20 °C in a sitting drop containing 0.2 μL of 18 mg/mL Rhs2CT-RhsI2-EF-Tu (25 mM MES-HCl 25 mM imidazole-HCl pH5.5, 150 mM NaCl, 1 mM MgCl_2_, 2 mM GDP and 0.5 mM TCEP) with 0.2 μL of reservoir solution (0.2 M ammonium acetate, 0.1 M HEPES pH 7.5 and 25% PEG3350). These crystals were used for microseeding to provide samples suitable for crystallographic analysis. Diffraction data were collected in-house using a Rigaku M007HF X-ray generator equipped with a Saturn 944HG+ CCD. The data were integrated as *P*12_1_1 with unit cell dimensions *a* 60.64 Å, *b* 107.04 Å, *c* 111.34 Å, β = 102.17°. A Matthews coefficient of 2.23 Å^3^ Da^−1^ suggested two heterotrimers in the asymmetric unit with solvent content of around 45% by volume. Subsequently, higher-resolution diffraction data were collected at Diamond Light Source (DLS; beamline I03), details in Supplementary Table 3. The structure of *E. coli* EF-Tu (PDB 1EFC) was used for molecular replacement calculations in PhaserMR^59^. The resulting electron density map at a resolution of 2.45 Å was of poor quality and the amino acid sequence for Rhs2CT-RhsI2 could not be reliably assigned to electron density.

To improve phase information for the Rhs2CT-RhsI2-EF-Tu complex, it was decided to determine the preliminary structure of RhsI2 separately. His_6_-tagged RhsI2 was purified and crystallised. Crystals appeared after a week in the condition containing 0.1 µL of 30 mg/ml RhsI2 (in 50 mM Tris pH 7.5, 250 mM NaCl) and 0.1 µL reservoir solution (Morpheus H1; Molecular Dimensions). These crystals were tested and 2.5 Å resolution data were collected in-house, as above. The data indicated space group *I*222 with unit cell dimensions *a* 67.64 Å, *b* 71.76 Å, *c* 78.05 Å. The model of RhsI2 generated in AlphaFold^26^ was used for molecular replacement calculations in PhaserMR. A Matthews coefficient of 2.25 Å^3^ Da^−1^ suggested two molecules in the asymmetric unit with solvent content of around 45% by volume. This preliminary structure of RhsI2 was partially refined (*R*_work_ = 0.31 and *R*_free_ = 0.35). Once the model has reached sufficient quality, it was used along with EF-Tu (PDB 1EFC) for molecular replacement calculations in PhaserMR. Following placement of the RhsI2 model, the electron density map improved, and it was possible to reliable include the third protein of the complex, Rhs2CT, guided by use of the AlphaFold model.

Rounds of electron and difference density map inspection, model manipulation in COOT ^60^ and refinement in REFMAC5^61^ led to a model consisting of two heterotrimers. *B*-factors were refined isotropically. Water molecules were assigned to well-defined peaks in the difference density map ( > 3.5 s) that were within 2.5–3.5 Å distance from hydrogen bond donor and acceptor groups. Mg^2+^ ions, molecules of 1,2-ethanediol, imidazole and ADP were also included. MolProbity^62^ was used in conjunction with the validation tools provided in COOT to assess model geometry throughout refinement. Crystallographic statistics are presented in Supplementary Table 3. The coordinates and structure factors have been deposited with the Protein Data Bank under accession code 9T38.

#### Rhs2CT-RhsI2 complex

An initial crystallisation condition was identified in the ProPlex screen (Molecular Dimensions; MD1-38) following four weeks incubation at 20 °C in a sitting drop condition containing 0.2 μL of 10 mg/mL Rhs2CT-RhsI2 (20 mM MES 20 mM imidazole pH 6.0, 150 mM NaCl, 1 mM MgCl_2_, 0.5 mM TCEP) and 0.2 μL of reservoir solution (0.2 M ammonium acetate, 0.1 M sodium acetate pH 5.0, 20% PEG4K). Crystals were optimised using microseeding, and diffraction data were collected at the European Synchrotron Radiation Facility (ESRF) on beamline ID23-1, equipped with a Dectris Pilatus3 X 2M detector. The data were integrated in the space group P 2_1_2_1_2_1_ with unit cell dimensions *a* 37.006 Å, *b* 81.931 Å, *c* 101.37 Å. A Matthews coefficient of 2.13 Å^3^ Da^−1^ suggested one heterodimer in the asymmetric unit with solvent content of around 40% by volume. The previously-solved structure of Rhs2CT-RhsI2-EF-Tu complex was used for molecular replacement calculations in PhaserMR. The resulting electron density map at a resolution of 2.17 Å was of good quality and refinement, water molecule assignment and model validation were carried out as described above. Acetate ions, a legacy of the crystallisation conditions, were included in the final model. Crystallographic statistics are presented in Supplementary Table 3. The coordinates and structure factors have been deposited in the Protein Data Bank under accession code 8CM0.

### Affinity isolation (‘pull-down’) of proteins interacting with Rhs2CT from total lysate samples

For preparation of bacterial lysates, cultures of *E. coli* MG1655 were grown in 25 ml LB at 37°C until an OD_600_ of 0.8–1.0. Cells were harvested by centrifugation and resuspended in 1 ml of lysis buffer containing 50 mM Tris-HCl (pH 7.5), 150 mM NaCl, 75 mM imidazole, 0.5% Triton X-100, protease inhibitor cocktail (Abcam, ab274282; 1:10,000), and DNase I (Sigma, 10104159001; ~1 mg per 10 ml). Lysis was performed by sonication (six pulses of 15 s at 30% amplitude, with 30 s intervals on ice) and lysates were clarified by centrifugation at 15,000 rpm for 20 min at 4°C. For preparation of fungal lysates, cells of *S. cerevisiae* K699 were grown to an OD_600_ of 1 in 25 ml YPDA at 30°C, harvested, resuspended in 300 µl lysis buffer (25 mM Tris-HCl pH 7.5, 150 mM NaCl, 0.1% Triton X-100, cOmplete EDTA-free Protease Inhibitor) and added to 300 µl of acid-washed glass beads. The cells were broken using a bead beater (5x 20 s at 5400 rpm, with 2 min interval on ice) and cell debris removed by centrifugation at 15000 rpm (21000 x*g*) for 10 min. For each individual pull-down, a 30 µl aliquot of HisPur™ Ni-NTA magnetic beads (ThermoFisher Scientific, Cat. No. 88832) was equilibrated by washing three times with wash buffer (50 mM Tris-HCl pH 7.5, 150 mM NaCl, 75 mM imidazole, 0.1% Triton X-100, 30 µM Zn(OAc)_2_, and 0.5 mM TCEP-HCl) and then incubated with 20 µg of purified His-tagged Rhs2CT protein in 1 ml of wash buffer for 1 h at 4°C on a tube rotator (40 rpm). The beads were washed three times with 1 ml of wash buffer to remove unbound protein and transferred to a fresh tube. Clarified bacterial or fungal lysates were then added to the protein-bound magnetic beads and incubated for 1 h at 4°C with rotation (40 rpm). Beads were then washed three times with wash buffer, transferred to a clean tube, and washed once more. Bound proteins were eluted by adding 30 µl of 2×SDS-PAGE sample buffer with 2.5% β-mercaptoethanol and incubating at 100°C for 5 min.

### Affinity isolation (‘pull-down’) of proteins interacting with Rhs2CT from *E. coli* cells

Cultures of *E. coli* BL21(DE3) carrying pET15b, pSC3251 or pSC3252 were grown in 25 ml LB + Ap for 3 h at 37°C to OD_600_ of 0.8 followed by induction with 10 µM IPTG for 2 h. Cells were recovered by centrifugation and resuspended in 1 ml of lysis buffer (50 mM Tris-HCl pH8.0, 150 mM NaCl, 20 mM imidazole, 0.5% Triton X100) containing cOmplete EDTA-free Protease Inhibitor (Cat No. 11836170001, Roche). Cell lysates were prepared by sonication (6x 15 s pulses with 30 s interval on ice) and clarified by centrifugation at 21,000 x*g* for 20 min at 4°C. The supernatant was incubated with His Pur^TM^ Ni-NTA magnetic beads previously washed three times in wash buffer (20 mM Tris-HCl pH 8.0, 100 mM NaCl, 50 mM imidazole and 0.1% Triton X100) for 1.5 h at 4°C. Following incubation, beads were washed three times in 1 ml wash buffer and then bound proteins were eluted using 30 μl of 2x SDS-PAGE sample buffer (100 mM Tris-HCl, pH 6.8, 3.2% SDS, 3.2 mM EDTA, 16% glycerol, 0.2 mg/ml bromophenol blue, 2.5% β-mercaptoethanol) and heated to 100°C for 5 min.

### Nuclease assay

To assess nuclease activity of refolded Rhs2CT *in vitro*, 1 µl of 1 mg/ml Rhs2CT or its catalytically-inactive variant, Rhs2CT_CM_, corresponding to a final protein concentration of approximately 3 µM, was added to a reaction mixture containing 1000 ng of plasmid DNA (pACYC-Duet) or *S. marcescens* total RNA in buffer *C2* and incubated at 37°C for 15 min. DNA was separated by 1% agarose gel electrophoresis and stained with GelRed (Biotium). RNA was separated by 10% TBE-Urea gel electrophoresis (Bio-Rad) and stained with SYBR-Gold^TM^ (Invitrogen).

### Co-culture assays for T6SS-mediated anti-fungal and anti-bacterial activity

Anti-fungal activity assays were based on the method reported previously^7^. The *S. marcescens* attacker strain and the fungal target strains *S. cerevisiae*, *C. glabrata* or *C. albicans* were normalised to an OD_600_ of 1 and mixed at a volume ratio of 1:1 (attacker:target cell ratio ~100:1) and 12.5 µl of the mixture were spotted on solid SC + 2% glucose solid media and incubated for 7 h at 30°C. Following incubation, the cells were harvested, serially diluted in YPDA and appropriate dilutions spread onto YPDA + Sm in order to enumerate the recovery of viable fungal cells. For the co-culture assay utilising *S. cerevisiae* expressing RhsI2, or a control construct, as the target strain, the co-culture, and pre-growth of the fungal target strains, were performed on SC-LEU + 2% raffinose with 1% galactose to induce expression of RhsI2.

Anti-bacterial activity assays were based on the method reported previously^63^. Attacker and target cells were normalised to an OD_600_ of 0.5 in LB, combined in a 1:1 ratio (v:v) and 25 µl of the mixture spotted on LB solid media. Co-culture assays were incubated for 4 h at 30°C for *S. marcescens* target and 37°C for *E. coli* target. Following incubation, the cells were harvested, serially diluted in LB and appropriate dilutions spread onto LB + Sm (JAD17 target) or LB + Kn (*E. coli* BW25113 target) in order to enumerate the recovery of viable target bacterial cells.

### Plate toxicity assay

For toxicity assays in yeast, cells of *S. cerevisiae* were pre-grown in non-inducing liquid minimal media (SC-LEU + 2% raffinose) at 30°C, adjusted to an OD_600_ of 1, serially diluted and 5 µl spotted onto SC-LEU + 2% raffinose agar containing galactose (0.04-1%) or glucose (1%) for the induction or repression, respectively, of gene expression. Agar plates were incubated at 30°C and pictures taken after 72-96 h when colonies of control strains had grown. When examining the phenotype of the conditional importin-α mutant *srp1-31*, *S. cerevisiae* strains were pre-grown in non-inducing liquid minimal media (SC-LEU + 2% raffinose) at 23°C for 36 h, adjusted to an OD_600_ of 1, serially diluted and 5 µl spotted onto minimal media agar as above. Agar plates were incubated at 30°C or 23°C, with pictures taken after 72-96 h or 6-7 days, respectively. For toxicity assays in bacteria, cells of *E. coli* MG1655 freshly transformed with pBAD18-Kn-derived plasmids were grown overnight on solid LB + Kn + 0.5% glucose media. adjusted to an OD_600_ of 1, serially diluted and 5 µl spotted onto M9 plates (1x M9 Salts, 2 mM MgSO_4_, 1 mM CaCl_2_, 0.5% glycerol) containing 0.002%-0.2% L-arabinose or 0.2% glucose for the induction or repression, respectively, of gene expression. Agar plates were incubated at 37°C and pictures taken after 24 h. Plate toxicity assays were repeated at least twice independently and representative images are shown.

### Immunoblot analysis

Cells of *S. cerevisiae* were grown to an OD_600_ of 0.8 at 30 °C in 25ml SC-LEU + 2% raffinose, followed by induction with 1% galactose for 3 h. Cells were harvested, resuspended in 300 µl lysis buffer (10 mM Tris-HCl pH 7.5, 150 mM NaCl, 0.1% Triton X-100, cOmplete EDTA-free Protease Inhibitor) and added to 300 µl of acid washed glass beads. The cells were broken using a bead beater (4x 20 s at 5400 rpm, with 2 min interval on ice) and cell debris removed by centrifugation at 14000 rpm (21000 x*g*) for 10 min. Cell lysate (total protein) samples were combined with 2x SDS-PAGE sample buffer (100 mM Tris-HCl, pH 6.8, 3.2% SDS, 3.2 mM EDTA, 16% glycerol, 0.2 mg/ml bromophenol blue, 2.5% β-mercaptoethanol) and boiled for 5 min. Proteins were separated by 4-20% SDS-PAGE and electroblotted onto polyvinylidine fluoride (PVDF, Millipore). GFP was detected by hybridisation of the primary antibody, monoclonal mouse anti-GFP (Roche #1814460001; 1:5,000) and peroxidase-conjugated anti-mouse secondary antibody (Bio-Rad #0170-6516; 1:10,000). Visualisation was performed an enhanced chemiluminescent detection kit (Millipore) and the Azure 600 imaging system (Cambridge Bioscience).

### Microscopy imaging of fungal cells

Cells of *S. cerevisiae* were grown at 30°C in YPDA supplemented with 2% glucose for 8 h, then washed twice in SC media by centrifugation at 1000 x*g*, resuspended in SC media, and used to inoculate a 5 ml culture in SC media containing 2% raffinose from a starting OD_600_ of 0.07. Following overnight growth, the cells were back-diluted to an OD_600_ of 0.2 and grown in 5 ml SC media containing 2% raffinose for 8 h, then used to inoculate 25 ml of SC media containing 2% raffinose to an OD_600_ of 0.07 and grown overnight. This overnight culture was used to inoculate 25 ml SC media containing 2% raffinose and 1% galactose to a starting OD_600_ of 0.5 and the culture was grown for 3 h to allow for induction of gene expression. Cells were then harvested by centrifugation at 1000 x*g* for 5 min. The cells were washed once in 1xPBS by centrifugation and then fixed in 4% PFA + 20% EtOH in 1xPBS overnight at 4°C. The cells were then washed with 1xPBS, incubated in 50 mmol NH_4_Cl for 5 minutes, washed with PBS again and spotted on an 18×18 mm No. 1.5H high precision glass coverslip (Marienfeld) coated in 0.1 mg/ml Poly-D-Lysine (PDL). After a 30 minute incubation, the coverslips were mounted onto slides with ProLong™ Glass Antifade Mountant with NucBlue™ Stain (Fisher P36981) and cured for 48 hours at room temperature to achieve the optimal refractive index prior to imaging. For experiments involving the temperature-sensitive strain NOY612, *S. cerevisiae* were grown at 23°C prior to induction; derivatives of K699 were then grown at 30°C for 3 hr in SC with 2% raffinose and 1% galactose, as above, while derivatives of NOY612 were grown at 30°C in SC with 2% raffinose for 30 min before the addition of 1% galactose and a further 3 h growth at 30°C. For quantifying DNA damage by TUNEL assay, Click-iT™ EdU Cell Proliferation Kit for Imaging with Alexa Fluor™ 647 dye (Fisher C10340) was used. Prior to the assay, yeast cells were fixed in 4% PFA + 20% EtOH in 1xPBS overnight at 4°C followed by digestion with 120 µg/ml Zymolyase 20T (Fisher 15445459) for 1 h at 37°C, followed by incubation in a permeabilization solution (2% Triton X-100 in 1x PBS). For bacterial cells, the procedure was the same except digestion with Zymolyase was omitted. The manufacturer’s protocol for the TUNEL assay on coverslips was then followed.

All imaging was performed on fixed cells mounted in ProLong™ Glass Antifade Mountant with NucBlue™ Stain (Fisher P36981). Imaging was performed on a Leica SP8 TCS SP8 3X STED microscope with a Leica HC PL APO 100x/1.40 Oil STED WHITE objective (Leica Microsystems). Samples were excited using a Supercontinuum White Light Laser at 488 nm, 580nm, 650 nm, and a Leica SP8 405 nm laser and detected using a PMT detector with the gain of 800 or Leica HyD hybrid detector with the gain of 100; detection windows were set to 415-460 nm for 405 nm activation; 500 – 550 nm for 488 nm activation, 610 – 660 nm for 580 nm activation and 660 – 730 nm for 640 nm activation. Images were acquired according to Nyquist sampling (in x, y and z) and with sequential acquisition unless otherwise specified. Three dimensional stacks of multiple fields of view were taken for each sample and experiments were repeated three times unless otherwise specified. TUNEL assay images were acquired using Nyquist sampling in x and y and 500 nm step in z.

All data, unless specified otherwise, was deconvolved using Huygens Professional software (SVI) with subsequent file formats maintaining metadata and bit depth. All analysis was performed on three-dimensional datasets with images in Figures 5 and 6 presented as single slices extracted from the three-dimensional volume. Representative maximum intensity projection images (all slices superimposed) are provided for EGFP and NucBlue channels in Supplementary Figure 5. Colocalisation analysis was performed using ImageJ Fiji with the colocalisation threshold plugin (https://imagej.net/plugins/colocalization-threshold) and automatic Costes threshold determination to give R coloc. For fluorescence intensity measurements, microscope acquisition parameters were maintained between samples. Thresholding was applied to select cells, or sub-cellular compartments, and mean intensity for the selected sub-regions measured using Fiji. One-way Analysis of Variance (ANOVA) with Šídák’s multiple comparison test was used for P value determination, and Standard Error of the Mean (SEM) for errors.

## Data availability statement

Coordinates and structure factors have been deposited with the Protein Data Bank under accession codes 9T38 (Rhs2CT-RhsI2-EF-Tu) and 8CM0 (Rhs2CT-RhsI2). All other data supporting the findings of this study are available within the paper and its supplementary information files.

## Acknowledgements

This work was supported by Wellcome (grant numbers 215599/Z/19/Z, S.J.C, W.N.H, J.Q. & C.R; 104556/Z/14/Z, Senior Research Fellowship S.J.C.; 220321/Z/20/Z, Senior Research Fellowship Renewal S.J.C.) and a Heriot-Watt James Watt PhD studentship (T.S.). The Diamond Synchrotron Light source (beamline I03) and European Synchrotron Radiation Facility (beamline ID23-1) are acknowledged for beamtime. We thank Grant Buchanan, Yi-Chia Liu, Laura Monlezun, Martin Hagan and Ramses Gallegos-Monterrosa for construction of strains and/or plasmids and contributing to preliminary work. We thank Christopher Earl for construction of vector pSC4135, Alessandra da Silva Dantas for generation of SN148 *mec1*Δ, and Paul Fyfe for assistance with crystallography. We thank Jessica Valli and the Edinburgh Super-Resolution Imaging Consortium (ESRIC) for access to advanced fluorescence microcopy and analysis, and Karim Labib and Kiran Madura for donation of yeast strains. For the purpose of Open Access, the authors have applied a CC BY public copyright licence to any Author Accepted Manuscript version arising from this submission.

## Author Contributions

S.J.C., G.M.A and G.P. conceived the study. G.M.A., G.P., and T.S. performed experimental work and analysed data. G.M.A., G.P., J.Q., W.N.H, C.R., and S.J.C designed experiments, analysed data and interpreted data. S.J.C, G.M.A., and G.P. wrote the manuscript with contributions and input from the other authors.

