## Supplementary Data for "An Rhs effector uses distinct target cell functions to intoxicate bacterial and fungal competitors"

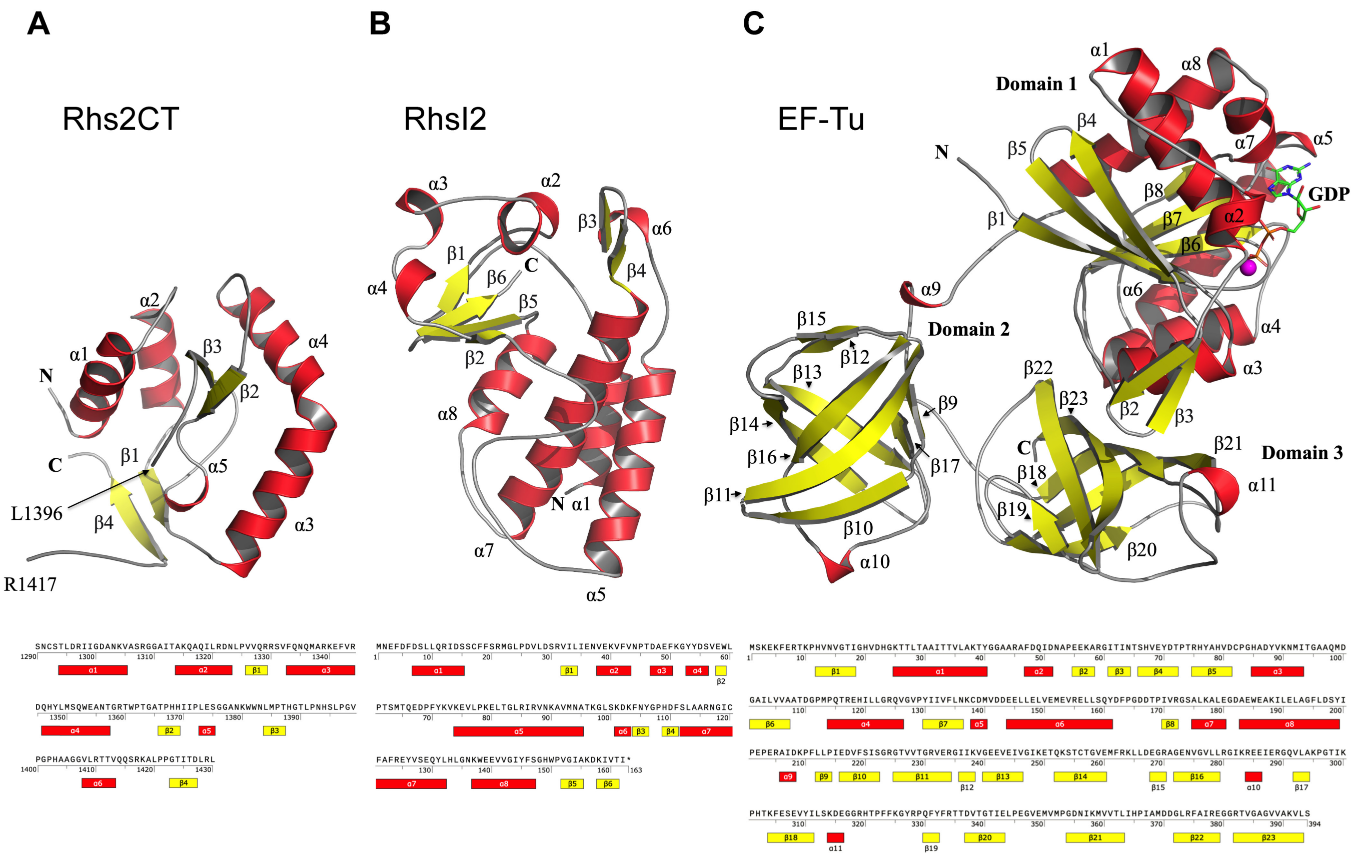

**Supplementary Figure 1. The structures of Rhs2CT, RhsI2 and EF-Tu.** Cartoon representation of the structures of **(A)** Rhs2CT from *S. marcescens* Db10, **(B)** RhsI2 from *S. marcescens* Db10 and **(C)** EF-Tu from *E. coli* BL21(DE3), each extracted from the heterotrimeric Rhs2CT-RhsI2-EF-Tu complex. Helices are shown in red, strands in yellow and loops in grey. Below each structure is the amino acid sequence showing elements of secondary structure assigned based on AlphaFold prediction and obtained structure.

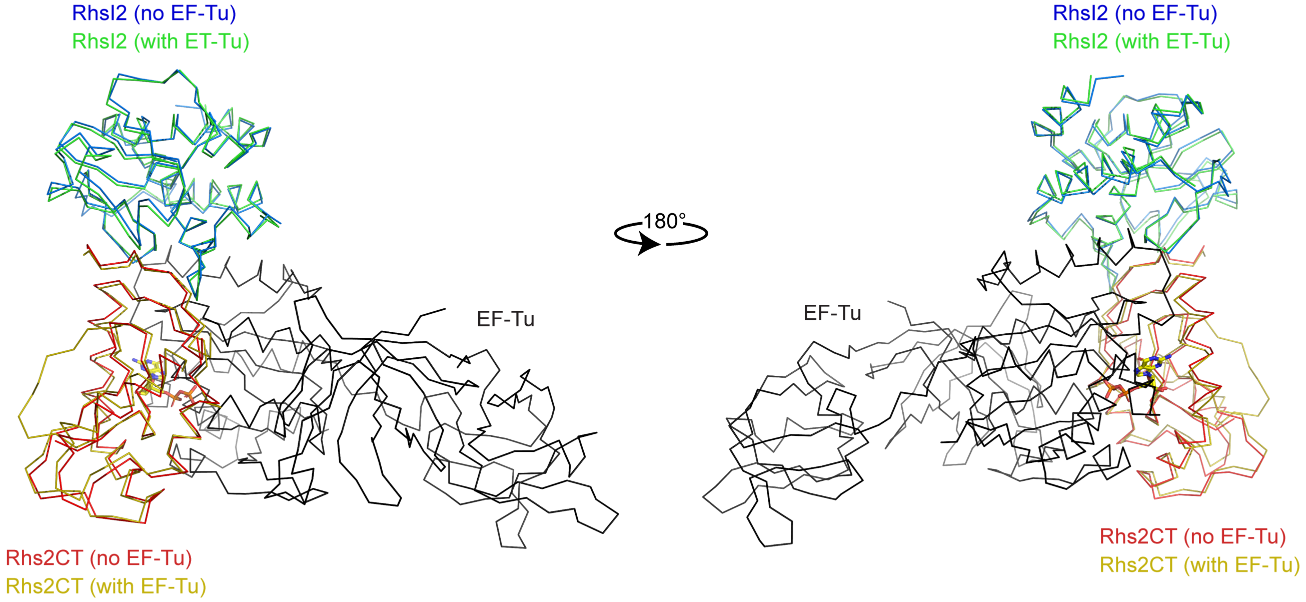

| Protein structure 1 (in Rhs2CT-RhsI2-EF-Tu heterotrimer) | Protein structure 2 (in Rhs2CT-RhsI2 heterodimer or EF-Tu apo) | RMSD | Number of residues aligned |
|---|---|---|---|
| Rhs2CT-RhsI2-EF-Tu | Rhs2CT-RhsI2 | 0.85 Å | 266 |
| Rhs2CT | Rhs2CT | 0.61 Å | 112 |
| RhsI2 | RhsI2 | 0.35 Å | 154 |
| EF-Tu | EF-Tu (PDB 1EFC) | 0.65 Å | 383 |

**Supplementary Figure 2. Complex formation with EF-Tu does not significantly change the structure of Rhs2CT, RhsI2 or the Rhs2CT-RhsI2 effector-immunity complex.** Top: overlay of the structure of the Rhs2CT-RhsI2 complex alone (Rhs2CT in red, RhsI2 in blue) or when in complex with EF-Tu (Rhs2CT in mustard, RhsI2 in green, EF-Tu in dark grey). Bottom: table of RMSD values obtained when the structures of Rhs2CT-RhsI2-EF-Tu, or constituent proteins within the heterotrimeric complex, are compared with those of the Rhs2CT-RhsI2 complex alone, constituent proteins within the heterodimeric complex, or EF-Tu alone (apo, PDB 1EFC).

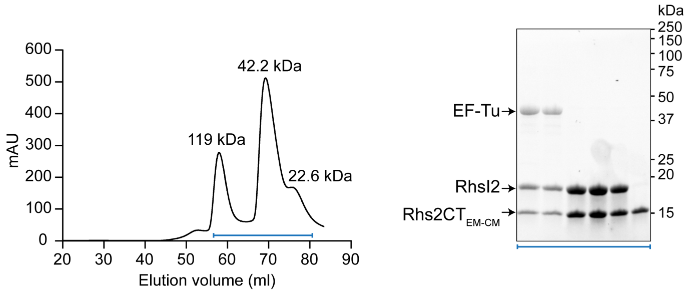

**Supplementary Figure 3. Purification of Rhs2CT$_{EM-CM}$-RhsI2 complex.** Size-exclusion chromatography using a 75 16/60 column of the protein complex isolated from *E. coli* producing recombinant His$_6$-Rhs2CT$_{EM-CM}$ and untagged RhsI2. Eluted proteins were visualised by SDS-PAGE and Coomassie staining (inset), with the fractions analysed indicated by the blue bracket, and the estimated molecular weight corresponding to the elution volume of each peak is noted.

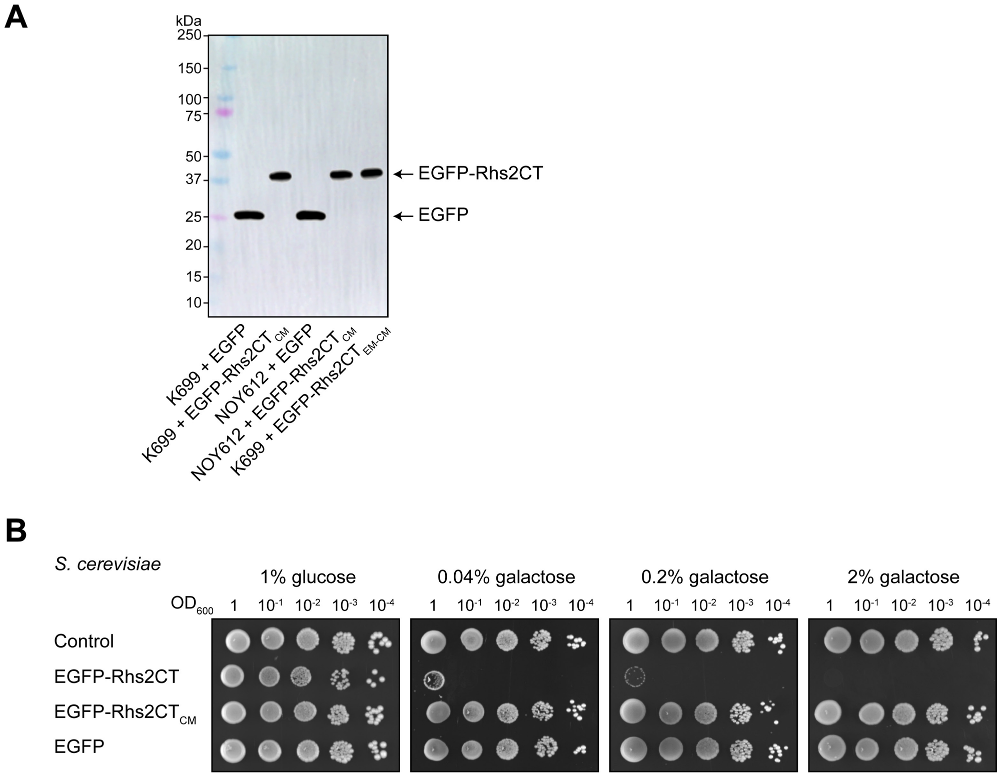

**Supplementary Figure 4. EGFP-Rhs2CT fusion proteins are stable and retain toxicity. (A)** Immunoblot detection of EGFP or EGFP-Rhs2CT in total protein samples of *S. cerevisiae* K699 or NOY612 chromosomal integration strains carrying constructs directing the expression of EGFP alone, EGFP-Rhs2CT$_{CM}$ or EGFP-Rhs2CT$_{CM-EM}$, as indicated, using an anti-GFP antibody. Expression was induced using 1% galactose. **(B)** Growth of *S. cerevisiae* K699 chromosomal integration strains carrying the empty promoter construct (control), or constructs directing the expression of EGFP alone or EGFP fused to the N-terminus of wild type Rhs2CT or of the catalytically inactive variant Rhs2CT$_{CM}$, on media containing glucose or galactose for repression or induction, respectively, of gene expression.

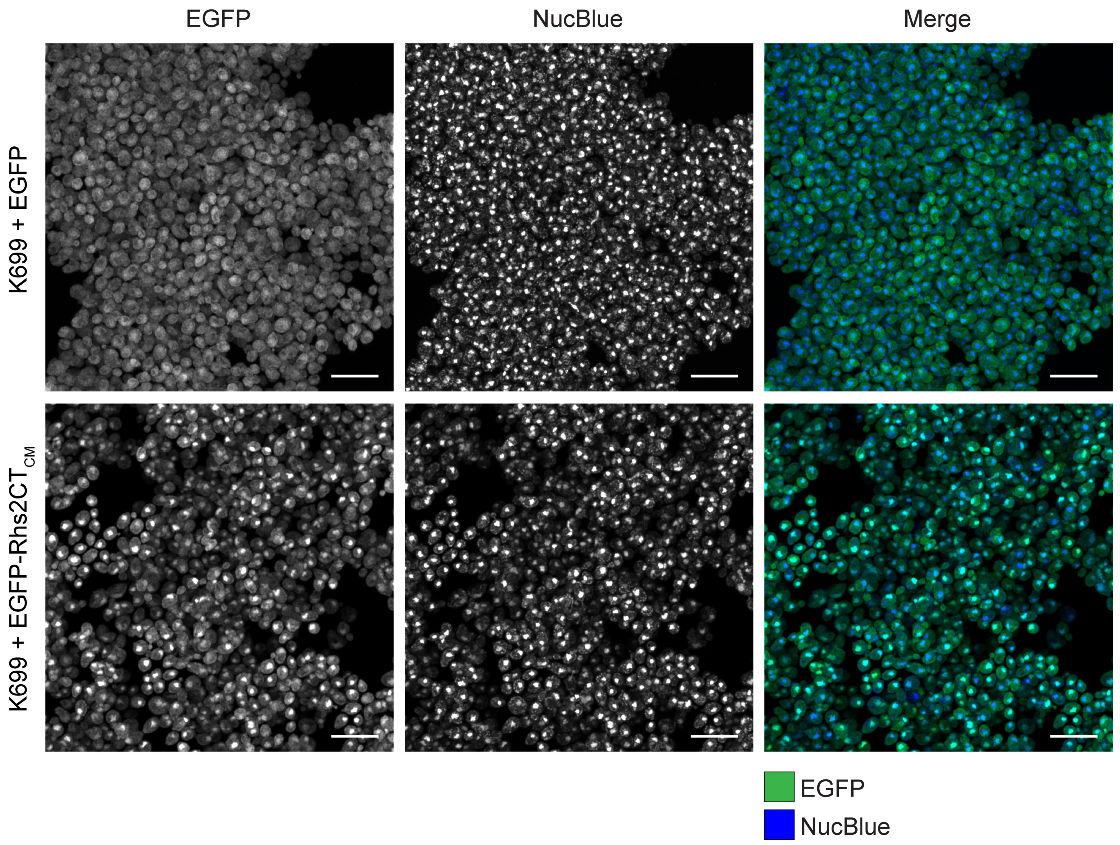

**Supplementary Figure 5. Maximum intensity projection images of *S. cerevisiae* expressing EGFP or EGFP-Rhs2CT$_{CM}$.** Representative maximum intensity projection images of *S. cerevisiae* K699 expressing EGFP alone or EGFP fused to the N-terminus of catalytically-inactive Rhs2CT (EGFP-Rhs2CT$_{CM}$, H1369A), shown as single channels of EGFP, NucBlue (nuclear stain) and the merged channels. Scale bars are 10 µm.

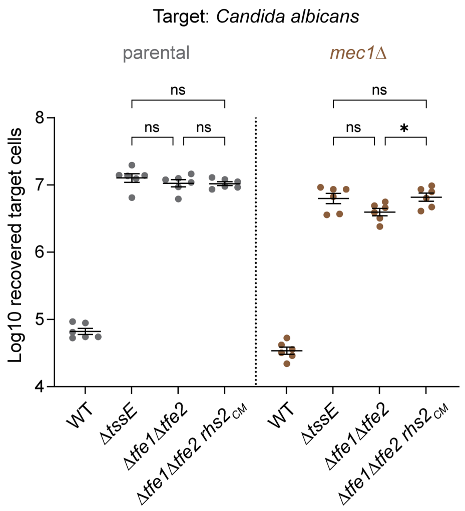

**Supplementary Figure 6. A mutant of *Candida albicans* with impaired nuclear DNA damage response becomes susceptible to T6SS-delivered Rhs2.** Recovery of *C. albicans* SN148 (parental) and the derivative lacking both alleles of *MEC1* (*mec1*Δ) following co-culture with wild type *S. marcescens* Db10 or mutants carrying in-frame gene deletions or point mutation in *rhs2* as indicated ($Rhs2_{CM}$, Rhs2 H1369A). Data are presented as mean ± SEM with individual data points overlaid (n=6 biological replicates; * $P<0.05$, ns not significant; one-way ANOVA with Šídák's test).

**Supplementary Table 1. Bacterial strains, fungal strains and plasmids used in this study.**

| Name | Description/genotype | Source / Reference |
|---|---|---|
| **Bacterial strains** | | |
| *Serratia marcescens* | | |
| Db10 | Wild type model strain | [1] |
| SJC11 | Db10 Δ*tssE* (Δ*SMDB11_2271*) | [2] |
| JAD13 | Db10 Δ*rhs2* (Δ*SMDB11_1610*) | [3] |
| JAD17 | Db10 Δ*rhs2* Δ*rhsI2* (Δ*SMDB11_1610-SMDB11_1611*) Sm$^R$ | [3] |
| FRC28 | Db10 Δ*vgrG1* Δ*rhs1* (Δ*SMBD11_2244, SMBD11_2278*) | [4] |
| FRC30 | Db10 Δ*vgrG1* Δ*rhs1* Δ*rhs2* | [4] |
| KT149 | Db10 Δ*tfe1* Δ*tfe2* (Δ*SMBD11_1112,* Δ*SMBD11_1083*) | [5] |
| YL44 | Db10 *rhs2*$_{CM}$. Encodes SMDB11_1610 H1369A (Rhs2CT$_{CM}$) at the normal chromosomal location. | This study |
| YL28 | Db10 Δ*vgrG1* Δ*rhs1* *rhs2*$_{CM}$ | This study |
| GMA18 | Db10 Δ*tfe1* Δ*tfe2* *rhs2*$_{CM}$ | This study |
| GB21 | Db10 *rhs2*$_{EM}$. Encodes SMDB11_1610 E1341R R1344A A1378W K1380E (Rhs2CT$_{EM}$) at the normal chromosomal location. | This study |
| GB20 | Db10 Δ*vgrG1* Δ*rhs1* *rhs2*$_{EM}$ | This study |
| GB35 | Db10 Δ*tfe1* Δ*tfe2* *rhs2*$_{EM}$ | This study |
| *Escherichia coli* | | |
| MG1655 | Wild type (model K-12 strain) | [6] |
| BL21(DE3) | Protein overexpression strain. Chromosomal λDE3 encodes IPTG-inducible T7 RNA polymerase. | Novagen |
| BW25113 *lacA*::Kn | Deletion of *lacA* with insertion of Kn-resistance cassette in BW25113, retrieved from the Keio mutant library; Kn$^R$ | [7] |
| CC118λ*pir* | Cloning host and donor strain for pKNG101-derived allelic exchange plasmids (λ*pir*) | [8] |
| HH26 pNJ5000 | Mobilizing strain for conjugal transfer | [9] |
| **Fungal strains** | | |
| *Saccharomyces cerevisiae* | | |
| K699 | *MAT***a** *ade2-1 trp1-1 leu2-3,112 HIS3-11,15 ura3 can1-100* | [10] |
| GMA10 | K699 with P$_{GAL10}$-P$_{GAL1}$ (promoter-only control) integrated in the *HIS3* locus (pSC3212). | This study |
| GMA07 | K699 P$_{GAL10}$-RhsI2. Construct directing the expression of RhsI2 (SMDB11_1611) under the control of the *GAL10* promoter integrated in the *HIS3* locus (pSC3233). | This study |
| GMA30 | K699 P$_{GAL1}$-Rhs2CT. Construct directing the expression of Rhs2CT (SMDB11_1610 amino acids 1290-1340) under the control of the *GAL1* promoter integrated in the *HIS3* locus (pSC3211). | This study |
| GMA37 | K699 P$_{GAL1}$-Rhs2CT$_{CM}$. Construct directing the expression of Rhs2CT$_{CM}$ (SMDB11_1610 amino acids 1290-1340, H1369A) under the control of the *GAL1* promoter integrated in the *HIS3* locus (pSC3213). | This study |
| GMA41 | K699 P$_{GAL10}$-RhsI2 P$_{GAL1}$-Rhs2CT. Construct directing the expression of Rhs2CT under the control of the *GAL1* promoter and RhsI2 under the control of the *GAL10* promoter integrated in the *HIS3* locus (pSC3204). | This study |
| GB17 | K699 P$_{GAL1}$-Rhs2CT$_{EM}$. Construct directing the expression of Rhs2CT$_{EM}$ (SMDB11_1610 amino acids 1290-1340, E1341R R1344A A1378W K1380E) under the control of the *GAL1* promoter integrated in the *HIS3* locus (pSC3720). | This study |

| | | |
|---|---|---|
| GMA24 | K699 $P_{GAL1}$-EGFP. Construct directing the expression of EGFP under the control of the *GAL1* promoter integrated in the *HIS3* locus (pSC3215). | This study |
| GMA27 | K699 $P_{GAL1}$-EGFP-Rhs2CT. Construct directing the expression of a fusion of EGFP to the N-terminus of Rhs2CT under the control of the *GAL1* promoter integrated in the *HIS3* locus (pSC3206). | This study |
| GMA25 | K699 $P_{GAL1}$-EGFP-Rhs2CT$_{CM}$. Construct directing the expression of a fusion of EGFP to the N-terminus of Rhs2CT$_{CM}$ under the control of the *GAL1* promoter integrated in the *HIS3* locus (pSC3214). | This study |
| GMA52 | K699 $P_{GAL1}$-EGFP-Rhs2CT$_{EM-CM}$. Construct directing the expression of a fusion of EGFP to the N-terminus of Rhs2CT$_{EM-CM}$ (SMDB11_1610 amino acids1290-1340, E1341R R1344A A1378W K1380E, H1369A) under the control of the *GAL1* promoter integrated in the *HIS3* locus (pSC3256) | This study |
| NOY612 | *MATα ura3-1 trp1Δ63 ade2-1 leu2-3,112 his3-11 srp1-31* (S116F) | [13] |
| GMA47 | NOY612 $P_{GAL1}$-Rhs2CT (pSC3211) | This study |
| GMA45 | NOY612 $P_{GAL1}$-EGFP (pSC3215) | This study |
| GMA46 | NOY612 $P_{GAL1}$-EGFP-Rhs2CT$_{CM}$ (pSC3214) | This study |
| W303-1a | *MAT***a** *ade2-1 ura3-1 his3-11,15 trp1-1 leu2-3,112 can1-100* | Labib lab |
| YGDP5 | W303-1a *mec1Δ sml1Δ* (*MAT***a** *ade2-1, ura3-1 his3-11,15 trp1-1 leu2-3,112 can1-100 sml1Δ::HIS3 mec1Δ::ADE2*) | Labib lab |
| GMA58 | W303-1a + pRS313 (*MAT***a** *ade2-1 ura3-1 his3-11,15 trp1-1 leu2-3,112 can1-100 HIS3*). Used as the 'parental' control strain for YGDP5. | This study |
| *Candida glabrata* | | |
| ATCC2001 | Wild type | ATCC |
| *Candida albicans* | | |
| JC747 | SN148 + CIp30 (*HIS1, ARG4, URA3*). Used as the 'parental' control for JC1549. | This study |
| JC1549 | SN148 *mec1Δ*. Genotype *mec1::HIS1 mec1::ARG4* +CIp10 (*URA3*). | This study |
| **Plasmids** | | |
| pRSFDuet-1 | Protein overexpression vector for the co-expression of two target genes. Each multiple cloning site (MCS) is preceded by a T7 promoter and the first site allows for fusion of an N-terminal His$_6$ tag (Kn$^R$) | Novagen |
| pET15b-TEV | Vector for protein overexpression under the control of the T7 promoter. Permits fusion of an His$_6$ tag followed by a TEV protease cleavage site to the N-terminus of the overexpressed protein (Ap$^R$) | [11] |
| pBAD18-Kn | Arabinose-inducible expression vector; gene of interest is cloned downstream of the $P_{ara}$ promoter (Kn$^R$) | [12] |
| pKNG101 | Suicide vector for allelic exchange (Sm$^R$, *sacBR, mobRK2, oriR6K*) (Sm$^R$) | [13] |
| pSC1384 | Plasmid carrying empty $P_{GAL10}$-$P_{GAL1}$ promoter construct, for integration into the *S. cerevisiae HIS3* locus (*LEU2*, Ap$^R$) | [5] |
| pRS313 | Replicative shuttle vector for *S. cerevisiae* supplying the *HIS3* gene under its native promoter | [14] |
| pSC672 | Coding sequence for RhsI2 (SMA1611) in pBAD18-Kn. Used to protect *E. coli* cloning host from Rhs2CT toxicity during construction of yeast expression vectors. | [3] |
| pSC3502 | Coding sequence for His$_6$-TEV-Rhs2CT (SMDB11_1610, amino acids 1290-1340) in MCS1 and RhsI2 (SMDB11_1611) in MCS2 of pRSFDuet-1, for production of His$_6$-TEV-Rhs2CT and untagged RhsI2. | This study |
| pSC3712 | Coding sequence for His$_6$-TEV-Rhs2CT$_{EM-CM}$ (SMDB11_1610, amino acids 1290-1340) in MCS1 and RhsI2 (SMDB11_1611) in MCS2 of pRSFDuet-1, for production of His$_6$-TEV-Rhs2CT and untagged RhsI2. | This study |
| pSC3251 | Coding sequence for Rhs2CT$_{CM}$ (SMDB11_1610 amino acids 1290-1340, H1369A) in pET15b-TEV, for production of His$_6$-TEV-Rhs2CT$_{CM}$. | This study |

| | | |
|---|---|---|
| pSC3252 | Coding sequence for Rhs2CT$_{EM-CM}$ (SMDB11_1610 amino acids1290-1340, E1341R R1344A A1378W K1380E, H1369A) in pET15b-TEV, for production of His$_6$-TEV-Rhs2CT$_{EM-CM}$. | This study |
| pSC955 | Coding sequence for RhsI2 (SMDB11_1611) in pACYCDuet-1, for production of His$_6$-RhsI2 | This study |
| pSC4135 | Derivative of pBAD18-Kn with repressed translational start site incorporated to reduce gene expression levels. | This study |
| pSC3729 | Coding sequence for Rhs2CT in pSC4135 | This study |
| pSC3731 | Coding sequence for Rhs2CT + RhsI2 in pSC4135 | This study |
| pSC3728 | Coding sequence for Rhs2CT$_{CM}$ in pSC4135 | This study |
| pSC3732 | Coding sequence for Rhs2CT$_{EM}$ in pSC4135 | This study |
| pSC3730 | Coding sequence for Rhs2CT$_{EM}$ + RhsI2 in pSC4135 | This study |
| pSC3212 | Vector for integration of cloned inserts under the control of the bidirectional *GAL1* and *GAL10* promoters in the *HIS3* locus of *S. cerevisiae*. Derived from pSC1384 by inclusion of several additional restriction sites. | This study |
| pSC3233 | Coding sequence for RhsI2 (SMDB11_1611) downstream of the *GAL10* promoter in pSC3212. | This study |
| pSC3211 | Coding sequence for Rhs2CT (SMDB11_1610 amino acids 1290-1340) downstream of the *GAL1* promoter in pSC3212. | This study |
| pSC3213 | Coding sequence for Rhs2CT$_{CM}$ (SMDB11_1610 amino acids 1290-1340, H1369A) downstream of the *GAL1* promoter in pSC3212. | This study |
| pSC3204 | Coding sequence for Rhs2CT and RhsI2 in pSC3212, downstream of the *GAL1* and *GAL10* promoters, respectively. | This study |
| pSC3720 | Coding sequence for Rhs2CT$_{EM}$ (SMDB11_1610 amino acids 1290-1340, E1341R R1344A A1378W K1380E) downstream of the *GAL1* promoter in pSC3212. | This study |
| pSC3215 | Coding sequence for EGFP downstream of the *GAL1* promoter in pSC3212. | This study |
| pSC3206 | Coding sequence for a fusion of EGFP to the N-terminus of Rhs2CT downstream of the *GAL1* promoter in pSC3212. | This study |
| pSC3214 | Coding sequence for a fusion of EGFP to the N-terminus of Rhs2CT$_{CM}$ downstream of the *GAL1* promoter in pSC3212. | This study |
| pSC3256 | Coding sequence for a fusion of EGFP to the N-terminus of Rhs2CT$_{EM-CM}$ (SMDB11_1610 amino acids1290-1340, E1341R R1344A A1378W K1380E, H1369A) downstream of the *GAL1* promoter in pSC3212. | This study |
| pSC1817 | pKNG101-derived allelic exchange plasmid for introduction of the *rhs2$_{CM}$* allele, encoding Rhs2 H1369A (SMDB11_1610 H1369A), at the normal chromosomal location in *S. marcescens* Db10. | This study |
| pSC3718 | pKNG101-derived allelic exchange plasmid for introduction of the *rhs2$_{EM}$* allele, encoding Rhs2 E1341R R1344A A1378W K1380E (SMDB11_1610 E1341R R1344A A1378W K1380E), at the normal chromosomal location in *S. marcescens* Db10. | This study |

### Supplementary Table 2. Sequences of cloned inserts in plasmids

| Plasmid | Sequence of cloned DNA (restriction sites used for cloning underlined) | Comment |
|---|---|---|
| pSC3502 | CCATGGGCAGCAGCCATCATCATCATCATCACAGCAGCGGCGAAAACCTGTATTTTCAGGGCG<br>GATCCAATTGTTCGACTCTTGACCGTATTATTGGTGATGCGAACAAAGTCGCTTCGCGTGGTG<br>GAGCTATAACAGCAAAACAAGCTCAGATACTGAGAGATAATTTACCGGTAGTTCAGAGACGGA<br>GTGTTTTCCAAAATCAGATGGCTCGCAAAGAGTTCGTCAGAGATCAGCATTATCTGATGAGTC<br>AGTGGGAAGCCAATACAGGTAGAACCTGGCCGACAGGAGCTACCCCGCACCACATAATCCCAC<br>TAGAAAGTGGGGGGGCGAATAAATGGTGGAATTTGATGCCTACTCATGGCACTTTGCCTAATC<br>ATTCTTTACCTGGTGTACCTGGCCCACATGCCGCCGGTGGAGTGCTTCGAACAACTGTTCAAC<br>AAAGCAGAAAAGCACTGCCTCCTGGAACTATAACTGATTTGAGATTGTGAAAGCTT | His$_6$-TEV-Rhs2CT in MCS1 |
| | CATATGAATGAATTTGACTTTGATAGTTTACTACAGCGCATCGATAGCTCATGCTTTTTCTCA<br>AGAATGGGGCTCCCAGATGTTCTGGATAGTCGCGTGATTTTAATTGAAAATGTTGAGAAGGTG<br>TTTGTTAATCCCACTGATGCAGAGTTTAAGGGATACTATGATAGTGTGGAATGGTTGCCAACA<br>TCAATGACTCAGGAGGATCCTTTCTACAAAGTAAAGGAGGTATTACCTAAGGAACTTACTGGT<br>TTACGTATTCGAGTAAACAAGGCTGTCATGAATGCAACAAAGGGATTGTCTAAGGATAAATTC<br>AATTACGGGCCTCATGACTTCAGCTTAGCAGCCAGAAATGGAATTTGTTTTGCGTTCAGAGAG<br>TATGTTTCTGAACAGTATCTTCATTTAGGGAATAAGTGGGAAGAAGTTGTGGGTATATACTTT<br>TCTGGTCACTGGCCTGTGGGTATAGCAAAAGATAAGATAGTTACCATTTAACTCGAG | RhsI2 in MCS2 |
| pSC3712 | CCATGGGCAGCAGCCATCATCATCATCATCACAGCAGCGGCGAAAACCTGTATTTTCAGGGCG<br>GATCCAATTGTTCGACTCTTGACCGTATTATTGGTGATGCGAACAAAGTCGCTTCGCGTGGTG<br>GAGCTATAACAGCAAAACAAGCTCAGATACTGAGAGATAATTTACCGGTAGTTCAGAGACGGA<br>GTGTTTTCCAAAATCAGATGGCTCGCAAACGGTTCGTCGCAGATCAGCATTATCTGATGAGTC<br>AGTGGGAAGCCAATACAGGTAGAACCTGGCCGACAGGAGCTACCCCGCACGCCATAATCCCAC<br>TAGAAAGTGGGGGGTGGAATGAATGGTGGAATTTGATGCCTACTCATGGCACTTTGCCTAATC<br>ATTCTTTACCTGGTGTACCTGGCCCACATGCCGCCGGTGGAGTGCTTCGAACAACTGTTCAAC<br>AAAGCAGAAAAGCACTGCCTCCTGGAACTATAACTGATTTGAGATTGTGAAAGCTT | His$_6$-TEV-Rhs2CT$_{EM\text{-}CM}$ in MCS1 |
| | CATATGAATGAATTTGACTTTGATAGTTTACTACAGCGCATCGATAGCTCATGCTTTTTCTCA<br>AGAATGGGGCTCCCAGATGTTCTGGATAGTCGCGTGATTTTAATTGAAAATGTTGAGAAGGTG<br>TTTGTTAATCCCACTGATGCAGAGTTTAAGGGATACTATGATAGTGTGGAATGGTTGCCAACA<br>TCAATGACTCAGGAGGATCCTTTCTACAAAGTAAAGGAGGTATTACCTAAGGAACTTACTGGT<br>TTACGTATTCGAGTAAACAAGGCTGTCATGAATGCAACAAAGGGATTGTCTAAGGATAAATTC<br>AATTACGGGCCTCATGACTTCAGCTTAGCAGCCAGAAATGGAATTTGTTTTGCGTTCAGAGAG<br>TATGTTTCTGAACAGTATCTTCATTTAGGGAATAAGTGGGAAGAAGTTGTGGGTATATACTTT<br>TCTGGTCACTGGCCTGTGGGTATAGCAAAAGATAAGATAGTTACCATTTAACTCGAG | RhsI2 in MCS2 |
| pSC3251 | CATATGTCAAGTAATTGTTCGACTCTTGACCGTATTATTGGTGATGCGAACAAAGTCGCTTCG<br>CGTGGTGGAGCTATAACAGCAAAACAAGCTCAGATACTGAGAGATAATTTACCGGTAGTTCAG<br>AGACGGAGTGTTTTCCAAAATCAGATGGCTCGCAAAGAGTTCGTCAGAGATCAGCATTATCTG<br>ATGAGTCAGTGGGAAGCCAATACAGGTAGAACCTGGCCGACAGGAGCTACCCCGCACGCGATA<br>ATCCCACTAGAAAGTGGGGGGGCGAATAAATGGTGGAATTTGATGCCTACTCATGGCACTTTG<br>CCTAATCATTCTTTACCTGGTGTACCTGGCCCACATGCCGCCGGTGGAGTGCTTCGAACAACT<br>GTTCAACAAAGCAGAAAAGCACTGCCTCCTGGAACTATAACTGATTTGAGATTGTGAGGATCC | |
| pSC3252 | CATATGTCAAGTAATTGTTCGACTCTTGACCGTATTATTGGTGATGCGAACAAAGTCGCTTCG<br>CGTGGTGGAGCTATAACAGCAAAACAAGCTCAGATACTGAGAGATAATTTACCGGTAGTTCAG<br>AGACGGAGTGTTTTCCAAAATCAGATGGCTCGCAAACGGTTCGTCGCAGATCAGCATTATCTG<br>ATGAGTCAGTGGGAAGCCAATACAGGTAGAACCTGGCCGACAGGAGCTACCCCGCACGCCATA<br>ATCCCACTAGAAAGTGGGGGGTGGAATGAATGGTGGAATTTGATGCCTACTCATGGCACTTTG<br>CCTAATCATTCTTTACCTGGTGTACCTGGCCCACATGCCGCCGGTGGAGTGCTTCGAACAACT<br>GTTCAACAAAGCAGAAAAGCACTGCCTCCTGGAACTATAACTGATTTGAGATTGTGACTCGAG | |
| pSC955 | GAATTCGAATGAATTTGACTTTGATAGTTTACTACAGCGCATCGATAGCTCATGCTT<br>TTTCTCAAGAATGGGGCTCCCAGATGTTCTGGATAGTCGCGTGATTTTAATTGAAAA<br>TGTTGAGAAGGTGTTTGTTAATCCCACTGATGCAGAGTTTAAGGGATACTATGATAG<br>TGTGGAATGGTTGCCAACATCAATGACTCAGGAGGATCCTTTCTACAAAGTAAAGGA<br>GGTATTACCTAAGGAACTTACTGGTTTACGTATTCGAGTAAACAAGGCTGTCATGAA<br>TGCAACAAAGGGATTGTCTAAGGATAAATTCAATTACGGGCCTCATGACTTCAGCTT<br>AGCAGCCAGAAATGGAATTTGTTTTGCGTTCAGAGAGTATGTTTCTGAACAGTATCT<br>TCATTTAGGGAATAAGTGGGAAGAAGTTGTGGGTATATACTTTTCTGGTCACTGGCC<br>TGTGGGTATAGCAAAAGATAAGATAGTTACCATTTAAAAGCTT | |
| pSC4135 | GAATTCGGTACCGGTGATACCAGCATCGTCTTGATGCCCTTGGCAGCACCCTGCTAAGGAGGC<br>AACCATATGACTAGTGTCGACCTGCAGGAGCTCGCATGC | Replaces EcoRI-SphI fragment |
| pSC3729 | CATATGAGTAATTGTTCGACTCTTGACCGTATTATTGGTGATGCGAACAAAGTCGCT<br>TCGCGTGGTGGAGCTATAACAGCAAAACAAGCTCAGATACTGAGAGATAATTTACCG | |

| | | |
|---|---|---|
| | GTAGTTCAGAGACGGAGTGTTTTCCAAAATCAGATGGCTCGCAAAGAGTTCGTCAGA GATCAGCATTATCTGATGAGTCAGTGGGAAGCCAATACAGGTAGAACCTGGCCGACA GGAGCTACCCCGCACCACATAATCCCACTAGAAAGTGGGGGGGCGAATAAATGGTGG AATTTGATGCCTACTCATGGCACTTTGCCTAATCATTCTTTACCTGGTGTACCTGGC CCACATGCCGCCGGTGGAGTGCTTCGAACAACTGTTCAACAAAGCAGAAAAGCACTG CCTCCTGGAACTATAACTGATTTGAGATTGTGAGTCGAC | |
| pSC3731 | CATATGAGTAATTGTTCGACTCTTGACCGTATTATTGGTGATGCGAACAAAGTCGCT TCGCGTGGTGGAGCTATAACAGCAAAACAAGCTCAGATACTGAGAGATAATTTACCG GTAGTTCAGAGACGGAGTGTTTTCCAAAATCAGATGGCTCGCAAAGAGTTCGTCAGA GATCAGCATTATCTGATGAGTCAGTGGGAAGCCAATACAGGTAGAACCTGGCCGACA GGAGCTACCCCGCACCACATAATCCCACTAGAAAGTGGGGGGGCGAATAAATGGTGG AATTTGATGCCTACTCATGGCACTTTGCCTAATCATTCTTTACCTGGTGTACCTGGC CCACATGCCGCCGGTGGAGTGCTTCGAACAACTGTTCAACAAAGCAGAAAAGCACTG CCTCCTGGAACTATAACTGATTTGAGATTGTGACATGAATGAATTTGACTTTGATAG TTTACTACAGCGCATCGATAGCTCATGCTTTTTCTCAAGAATGGGGCTCCCAGATGT TCTGGATAGTCGCGTGATTTTAATTGAAAATGTTGAGAAGGTGTTTGTTAATCCCAC TGATGCAGAGTTTAAGGGATACTATGATAGTGTGGAATGGTTGCCAACATCAATGAC TCAGGAGGATCCTTTCTACAAAGTAAAGGAGGTATTACCTAAGGAACTTACTGGTTT ACGTATTCGAGTAAACAAGGCTGTCATGAATGCAACAAAGGGATTGTCTAAGGATAA ATTCAATTACGGGCCTCATGACTTCAGCTTAGCAGCCAGAAATGGAATTTGTTTTGC GTTCAGAGAGTATGTTTCTGAACAGTATCTTCATTTAGGGAATAAGTGGGAAGAAGT TGTGGGTATATACTTTTCTGGTCACTGGCCTGTGGGTATAGCAAAAGATAAGATAGT TACCATTTAAGTCGAC | |
| pSC3728 | CATATGAGTAATTGTTCGACTCTTGACCGTATTATTGGTGATGCGAACAAAGTCGCT TCGCGTGGTGGAGCTATAACAGCAAAACAAGCTCAGATACTGAGAGATAATTTACCG GTAGTTCAGAGACGGAGTGTTTTCCAAAATCAGATGGCTCGCAAAGAGTTCGTCAGA GATCAGCATTATCTGATGAGTCAGTGGGAAGCCAATACAGGTAGAACCTGGCCGACA GGAGCTACCCCGCACGCGATAATCCCACTAGAAAGTGGGGGGGCGAATAAATGGTGG AATTTGATGCCTACTCATGGCACTTTGCCTAATCATTCTTTACCTGGTGTACCTGGC CCACATGCCGCCGGTGGAGTGCTTCGAACAACTGTTCAACAAAGCAGAAAAGCACTG CCTCCTGGAACTATAACTGATTTGAGATTGTGAGTCGAC | |
| pSC3732 | CATATGAGTAATTGTTCGACTCTTGACCGTATTATTGGTGATGCGAACAAAGTCGCT TCGCGTGGTGGAGCTATAACAGCAAAACAAGCTCAGATACTGAGAGATAATTTACCG GTAGTTCAGAGACGGAGTGTTTTCCAAAATCAGATGGCTCGCAAACGGTTCGTCGCA GATCAGCATTATCTGATGAGTCAGTGGGAAGCCAATACAGGTAGAACCTGGCCGACA GGAGCTACCCCGCACCACATAATCCCACTAGAAAGTGGGGGGTGGAATGAATGGTGG AATTTGATGCCTACTCATGGCACTTTGCCTAATCATTCTTTACCTGGTGTACCTGGC CCACATGCCGCCGGTGGAGTGCTTCGAACAACTGTTCAACAAAGCAGAAAAGCACTG CCTCCTGGAACTATAACTGATTTGAGATTGTGAGTCGAC | |
| pSC3730 | CATATGAGTAATTGTTCGACTCTTGACCGTATTATTGGTGATGCGAACAAAGTCGCT TCGCGTGGTGGAGCTATAACAGCAAAACAAGCTCAGATACTGAGAGATAATTTACCG GTAGTTCAGAGACGGAGTGTTTTCCAAAATCAGATGGCTCGCAAACGGTTCGTCGCA GATCAGCATTATCTGATGAGTCAGTGGGAAGCCAATACAGGTAGAACCTGGCCGACA GGAGCTACCCCGCACCACATAATCCCACTAGAAAGTGGGGGGTGGAATGAATGGTGG AATTTGATGCCTACTCATGGCACTTTGCCTAATCATTCTTTACCTGGTGTACCTGGC CCACATGCCGCCGGTGGAGTGCTTCGAACAACTGTTCAACAAAGCAGAAAAGCACTG CCTCCTGGAACTATAACTGATTTGAGATTGTGACATGAATGAATTTGACTTTGATAG TTTACTACAGCGCATCGATAGCTCATGCTTTTTCTCAAGAATGGGGCTCCCAGATGT TCTGGATAGTCGCGTGATTTTAATTGAAAATGTTGAGAAGGTGTTTGTTAATCCCAC TGATGCAGAGTTTAAGGGATACTATGATAGTGTGGAATGGTTGCCAACATCAATGAC TCAGGAGGATCCTTTCTACAAAGTAAAGGAGGTATTACCTAAGGAACTTACTGGTTT ACGTATTCGAGTAAACAAGGCTGTCATGAATGCAACAAAGGGATTGTCTAAGGATAA ATTCAATTACGGGCCTCATGACTTCAGCTTAGCAGCCAGAAATGGAATTTGTTTTGC GTTCAGAGAGTATGTTTCTGAACAGTATCTTCATTTAGGGAATAAGTGGGAAGAAGT TGTGGGTATATACTTTTCTGGTCACTGGCCTGTGGGTATAGCAAAAGATAAGATAGT TACCATTTAAGTCGAC | |
| pSC3212 | GTCGACGGCTCCCCGGGTTAATTAAGGCGCGCCAGATCTCCTTGAATTTTCAAAAAT TCTTACTTTTTTTTTGGATGGACGCAAAGAAGTTTAATAATCATATTACATGGCATT ACCACCATATACATATCCATATACATATCCATATCTAATCTTACTTATATGTTGTGG | Replaces SalI-SphI fragment |

| | |
|---|---|
| | AAATGTAAAGAGCCCCATTATCTTAGCCTAAAAAAACCTTCTCTTTGGAACTTTCAG<br>TAATACGCTTAACTGCTCATTGCTATATTGAAGTACGGATTAGAAGCCGCCGAGCGG<br>GTGACAGCCCTCCGAAGGAAGACTCTCCTCCGTGCGTCCTCGTCTTCACCGGTCGCG<br>TTCCTGAAACGCAGATGTGCCTCGCGCCGCACTGCTCCGAACAATAAAGATTCTACA<br>ATACTAGCTTTTATGGTTATGAAGAGGAAAAATTGGCAGTAACCTGGCCCCACAAAC<br>CTTCAAATGAACGAATCAAATTAACAACCATAGGATGATAATGCGATTAGTTTTTTA<br>GCCTTATTTCTGGGGTAATTAATCAGCGAAGCGATGATTTTTGATCTATTAACAGAT<br>ATATAAATGCAAAAACTGCATAACCACTTTAACTAATACTTTCAACATTTTCGGTTT<br>GTATTACTTCTTATTCAAATGTAATAAAGTATCAACAAAAAATTGTTAATATACCT<br>CTATACTTTAACGTCAAGGAGAAAAAACCCCGGATCCTAATATTCTAGATACCTTCT<br>CGAGGCCTTTGCATGC |
| pSC3233 | AGATCTATGAATGAATTTGACTTTGATAGTTTACTACAGCGCATCGATAGCTCATGC<br>TTTTTCTCAAGAATGGGGCTCCCAGATGTTCTGGATAGTCGCGTGATTTTAATTGAA<br>AATGTTGAGAAGGTGTTTGTTAATCCCACTGATGCAGAGTTTAAGGGATACTATGAT<br>AGTGTGGAATGGTTGCCAACATCAATGACTCAGGAGGATCCTTTCTACAAAGTAAAG<br>GAGGTATTACCTAAGGAACTTACTGGTTTACGTATTCGAGTAAACAAGGCTGTCATG<br>AATGCAACAAAGGGATTGTCTAAGGATAAATTCAATTACGGGCCTCATGACTTCAGC<br>TTAGCAGCCAGAAATGGAATTTGTTTTGCGTTCAGAGAGTATGTTTCTGAACAGTAT<br>CTTCATTTAGGGAATAAGTGGGAAGAAGTTGTGGGTATATACTTTTCTGGTCACTGG<br>CCTGTGGGTATAGCAAAAGATAAGATAGTTACCATTTAAGCATGCGTCGAC |
| pSC3211 | GGATCCATGAGTAATTGTTCGACTCTTGACCGTATTATTGGTGATGCGAACAAAGTC<br>GCTTCGCGTGGTGGAGCTATAACAGCAAAACAAGCTCAGATACTGAGAGATAATTTA<br>CCGGTAGTTCAGAGACGGAGTGTTTTCCAAAATCAGATGGCTCGCAAAGAGTTCGTC<br>AGAGATCAGCATTATCTGATGAGTCAGTGGGAAGCCAATACAGGTAGAACCTGGCCG<br>ACAGGAGCTACCCCGCACCACATAATCCCACTAGAAAGTGGGGGGGCGAATAAATGG<br>TGGAATTTGATGCCTACTCATGGCACTTTGCCTAATCATTCTTTACCTGGTGTACCT<br>GGCCCACATGCCGCCGGTGGAGTGCTTCGAACAACTGTTCAACAAAGCAGAAAAGCA<br>CTGCCTCCTGGAACTATAACTGATTTGAGATTGTGATCTAGA |
| pSC3213 | GGATCCATGAGTAATTGTTCGACTCTTGACCGTATTATTGGTGATGCGAACAAAGTC<br>GCTTCGCGTGGTGGAGCTATAACAGCAAAACAAGCTCAGATACTGAGAGATAATTTA<br>CCGGTAGTTCAGAGACGGAGTGTTTTCCAAAATCAGATGGCTCGCAAAGAGTTCGTC<br>AGAGATCAGCATTATCTGATGAGTCAGTGGGAAGCCAATACAGGTAGAACCTGGCCG<br>ACAGGAGCTACCCCGCACGCGATAATCCCACTAGAAAGTGGGGGGGCGAATAAATGG<br>TGGAATTTGATGCCTACTCATGGCACTTTGCCTAATCATTCTTTACCTGGTGTACCT<br>GGCCCACATGCCGCCGGTGGAGTGCTTCGAACAACTGTTCAACAAAGCAGAAAAGCA<br>CTGCCTCCTGGAACTATAACTGATTTGAGATTGTGATCTAGA |
| pSC3204 | As for pSC3233 and pSC3211 |
| pSC3720 | GGATCCATGAGTAATTGTTCGACTCTTGACCGTATTATTGGTGATGCGAACAAAGTC<br>GCTTCGCGTGGTGGAGCTATAACAGCAAAACAAGCTCAGATACTGAGAGATAATTTA<br>CCGGTAGTTCAGAGACGGAGTGTTTTCCAAAATCAGATGGCTCGCAAACGGTTCGTC<br>GCAGATCAGCATTATCTGATGAGTCAGTGGGAAGCCAATACAGGTAGAACCTGGCCG<br>ACAGGAGCTACCCCGCACCACATAATCCCACTAGAAAGTGGGGGGTGGAATGAATGG<br>TGGAATTTGATGCCTACTCATGGCACTTTGCCTAATCATTCTTTACCTGGTGTACCT<br>GGCCCACATGCCGCCGGTGGAGTGCTTCGAACAACTGTTCAACAAAGCAGAAAAGCA<br>CTGCCTCCTGGAACTATAACTGATTTGAGATTGTGATCTAGA |
| pSC3215 | GGATCCTATACCATGGTGAGCAAGGGCGAGGAGCTGTTCACCGGGGTGGTGCCCATC<br>CTGGTCGAGCTGGACGGCGACGTAAACGGCCACAAGTTCAGCGTGTCCGGCGAGGGC<br>GAGGGCGATGCCACCTACGGCAAGCTGACCCTGAAGTTCATCTGCACCACCGGCAAG<br>CTGCCCGTGCCCTGGCCCACCCTCGTGACCACCCTGACCTACGGCGTGCAGTGCTTC<br>AGCCGCTACCCCGACCACATGAAGCAGCACGACTTCTTCAAGTCCGCCATGCCCGAA<br>GGCTACGTCCAGGAGCGCACCATCTTCTTCAAGGACGACGGCAACTACAAGACCCGC<br>GCCGAGGTGAAGTTCGAGGGCGACACCCTGGTGAACCGCATCGAGCTGAAGGGCATC<br>GACTTCAAGGAGGACGGCAACATCCTGGGGCACAAGCTGGAGTACAACTACAACAGC<br>CACAACGTCTATATCATGGCCGACAAGCAGAAGAACGGCATCAAGGTGAACTTCAAG<br>ATCCGCCACAACATCGAGGACGGCAGCGTGCAGCTCGCCGACCACTACCAGCAGAAC<br>ACCCCCATCGGCGACGGCCCCGTGCTGCTGCCCGACAACCACTACCTGAGCACCCAG<br>TCCGCCCTGAGCAAAGACCCCAACGAGAAGCGCGATCACATGGTCCTGCTGGAGTTC<br>GTGACCGCCGCCGGGATCACTCTCGGCATGGACGAGCTGTACAAGTAATCTAGA |

| | |
|---|---|
| pSC3206 | GGATCCTATACCATGGTGAGCAAGGGCGAGGAGCTGTTCACCGGGGTGGTGCCCATC<br>CTGGTCGAGCTGGACGGCGACGTAAACGGCCACAAGTTCAGCGTGTCCGGCGAGGGC<br>GAGGGCGATGCCACCTACGGCAAGCTGACCCTGAAGTTCATCTGCACCACCGGCAAG<br>CTGCCCGTGCCCTGGCCCACCCTCGTGACCACCCTGACCTACGGCGTGCAGTGCTTC<br>AGCCGCTACCCCGACCACATGAAGCAGCACGACTTCTTCAAGTCCGCCATGCCCGAA<br>GGCTACGTCCAGGAGCGCACCATCTTCTTCAAGGACGACGGCAACTACAAGACCCGC<br>GCCGAGGTGAAGTTCGAGGGCGACACCCTGGTGAACCGCATCGAGCTGAAGGGCATC<br>GACTTCAAGGAGGACGGCAACATCCTGGGGCACAAGCTGGAGTACAACTACAACAGC<br>CACAACGTCTATATCATGGCCGACAAGCAGAAGAACGGCATCAAGGTGAACTTCAAG<br>ATCCGCCACAACATCGAGGACGGCAGCGTGCAGCTCGCCGACCACTACCAGCAGAAC<br>ACCCCCATCGGCGACGGCCCCGTGCTGCTGCCCGACAACCACTACCTGAGCACCCAG<br>TCCGCCCTGAGCAAAGACCCCAACGAGAAGCGCGATCACATGGTCCTGCTGGAGTTC<br>GTGACCGCCGCCGGGATCACTCTCGGCATGGACGAGCTGTACAAGGGAGCAGGAGCA<br>CCGGTCGCCACCAGTAATTGTTCGACTCTTGACCGTATTATTGGTGATGCGAACAAA<br>GTCGCTTCGCGTGGTGGAGCTATAACAGCAAAACAAGCTCAGATACTGAGAGATAAT<br>TTACCGGTAGTTCAGAGACGGAGTGTTTTCCAAAATCAGATGGCTCGCAAAGAGTTC<br>GTCAGAGATCAGCATTATCTGATGAGTCAGTGGGAAGCCAATACAGGTAGAACCTGG<br>CCGACAGGAGCTACCCCGCACCACATAATCCCACTAGAAAGTGGGGGGGCGAATAAA<br>TGGTGGAATTTGATGCCTACTCATGGCACTTTGCCTAATCATTCTTTACCTGGTGTA<br>CCTGGCCCACATGCCGCCGGTGGAGTGCTTCGAACAACTGTTCAACAAAGCAGAAAA<br>GCACTGCCTCCTGGAACTATAACTGATTTGAGATTGTGATCTAGA |
| pSC3214 | GGATCCTATACCATGGTGAGCAAGGGCGAGGAGCTGTTCACCGGGGTGGTGCCCATC<br>CTGGTCGAGCTGGACGGCGACGTAAACGGCCACAAGTTCAGCGTGTCCGGCGAGGGC<br>GAGGGCGATGCCACCTACGGCAAGCTGACCCTGAAGTTCATCTGCACCACCGGCAAG<br>CTGCCCGTGCCCTGGCCCACCCTCGTGACCACCCTGACCTACGGCGTGCAGTGCTTC<br>AGCCGCTACCCCGACCACATGAAGCAGCACGACTTCTTCAAGTCCGCCATGCCCGAA<br>GGCTACGTCCAGGAGCGCACCATCTTCTTCAAGGACGACGGCAACTACAAGACCCGC<br>GCCGAGGTGAAGTTCGAGGGCGACACCCTGGTGAACCGCATCGAGCTGAAGGGCATC<br>GACTTCAAGGAGGACGGCAACATCCTGGGGCACAAGCTGGAGTACAACTACAACAGC<br>CACAACGTCTATATCATGGCCGACAAGCAGAAGAACGGCATCAAGGTGAACTTCAAG<br>ATCCGCCACAACATCGAGGACGGCAGCGTGCAGCTCGCCGACCACTACCAGCAGAAC<br>ACCCCCATCGGCGACGGCCCCGTGCTGCTGCCCGACAACCACTACCTGAGCACCCAG<br>TCCGCCCTGAGCAAAGACCCCAACGAGAAGCGCGATCACATGGTCCTGCTGGAGTTC<br>GTGACCGCCGCCGGGATCACTCTCGGCATGGACGAGCTGTACAAGGGAGCAGGAGCA<br>CCGGTCGCCACCAGTAATTGTTCGACTCTTGACCGTATTATTGGTGATGCGAACAAA<br>GTCGCTTCGCGTGGTGGAGCTATAACAGCAAAACAAGCTCAGATACTGAGAGATAAT<br>TTACCGGTAGTTCAGAGACGGAGTGTTTTCCAAAATCAGATGGCTCGCAAAGAGTTC<br>GTCAGAGATCAGCATTATCTGATGAGTCAGTGGGAAGCCAATACAGGTAGAACCTGG<br>CCGACAGGAGCTACCCCGCACGCGATAATCCCACTAGAAAGTGGGGGGGCGAATAAA<br>TGGTGGAATTTGATGCCTACTCATGGCACTTTGCCTAATCATTCTTTACCTGGTGTA<br>CCTGGCCCACATGCCGCCGGTGGAGTGCTTCGAACAACTGTTCAACAAAGCAGAAAA<br>GCACTGCCTCCTGGAACTATAACTGATTTGAGATTGTGATCTAGA |
| pSC3256 | GGATCCTATACCATGGTGAGCAAGGGCGAGGAGCTGTTCACCGGGGTGGTGCCCATCCTGGTC<br>GAGCTGGACGGCGACGTAAACGGCCACAAGTTCAGCGTGTCCGGCGAGGGCGAGGGCGATGCC<br>ACCTACGGCAAGCTGACCCTGAAGTTCATCTGCACCACCGGCAAGCTGCCCGTGCCCTGGCCC<br>ACCCTCGTGACCACCCTGACCTACGGCGTGCAGTGCTTCAGCCGCTACCCCGACCACATGAAG<br>CAGCACGACTTCTTCAAGTCCGCCATGCCCGAAGGCTACGTCCAGGAGCGCACCATCTTCTTC<br>AAGGACGACGGCAACTACAAGACCCGCGCCGAGGTGAAGTTCGAGGGCGACACCCTGGTGAAC<br>CGCATCGAGCTGAAGGGCATCGACTTCAAGGAGGACGGCAACATCCTGGGGCACAAGCTGGAG<br>TACAACTACAACAGCCACAACGTCTATATCATGGCCGACAAGCAGAAGAACGGCATCAAGGTG<br>AACTTCAAGATCCGCCACAACATCGAGGACGGCAGCGTGCAGCTCGCCGACCACTACCAGCAG<br>AACACCCCCATCGGCGACGGCCCCGTGCTGCTGCCCGACAACCACTACCTGAGCACCCAGTCC<br>GCCCTGAGCAAAGACCCCAACGAGAAGCGCGATCACATGGTCCTGCTGGAGTTCGTGACCGCC<br>GCCGGGATCACTCTCGGCATGGACGAGCTGTACAAGGGAGCAGGAGCACCGGTCGCCACCAGT<br>AATTGTTCGACTCTTGACCGTATTATTGGTGATGCGAACAAAGTCGCTTCGCGTGGTGGAGCT<br>ATAACAGCAAAACAAGCTCAGATACTGAGAGATAATTTACCGGTAGTTCAGAGACGGAGTGTT<br>TTCCAAAATCAGATGGCTCGCAAACGGTTCGTCGCAGATCAGCATTATCTGATGAGTCAGTGG<br>GAAGCCAATACAGGTAGAACCTGGCCGACAGGAGCTACCCCGCACGCCATAATCCCACTAGAA<br>AGTGGGGGGTGGAATGAATGGTGGAATTTGATGCCTACTCATGGCACTTTGCCTAATCATTCT<br>TTACCTGGTGTACCTGGCCCACATGCCGCCGGTGGAGTGCTTCGAACAACTGTTCAACAAAGC<br>AGAAAAGCACTGCCTCCTGGAACTATAACTGATTTGAGATTGTGATCTAGA |

| | |
|---|---|
| pSC1817 | TCTAGAATCTGAACAGCGCGCCGCTGGAGGTGACCGACGCGGCGGGCAACCTGTGCTGGTCCG<br>GGCAATACGACACCTTCGGCAAGCTGCAGGGCCAGACGGTGGCCGGCGCGGCGAAGCGGCAGG<br>GCGCGCAATACCAGCAGCCGCTGCGCTACGCCGGGCAATACCAGGATGACGAAAGTGGCCTGC<br>ACTACAACCTGTTCCGCTACTACGAACCCGAGGTGGGGCGTTTCACCACGCAGGATCCGATAG<br>GGTTGGAAGGCGGGCTGAATTTATATGCTTATGGGCCAAATCCTTTAACTTGGATAGATCCAT<br>TTGGGTTAAGTAATTGTTCGACTCTTGACCGTATTATTGGTGATGCGAACAAAGTCGCTTCGC<br>GTGGTGGAGCTATAACAGCAAAACAAGCTCAGATACTGAGAGATAATTTACCGGTAGTTCAGA<br>GACGGAGTGTTTTCCAAAATCAGATGGCTCGCAAAGAGTTCGTCAGAGATCAGCATTATCTGA<br>TGAGTCAGTGGGAAGCCAATACAGGTAGAACCTGGCCGACAGGAGGTACCCCGCACGCCATAA<br>TCCCACTAGAAAGTGGGGGGGCGAATAAATGGTGGAATTTGATGCCTACTCATGGCACTTTGC<br>CTAATCATTCTTTACCTGGTGTACCTGGCCCACATGCCGCCGGTGGAGTGCTTCGAACAACTG<br>TTCAACAAAGCAGAAAAGCACTGCCTCCTGGAACTATAACTGATTTGAGATTGTGACATGAAT<br>GAATTTGACTTTGATAGTTTACTACAGCGCATCGATAGCTCATGCTTTTTCTCAAGAATGGGG<br>CTCCCAGATGTTCTGGATAGTCGCGTGATTTTAATTGAAAATGTTGAGAAGGTGTTTGTTAAT<br>CCCACTGATGCAGAGTTTAAGGGATACTATGATAGTGTGGAATGGTTGCCAACATCAATGACT<br>CAGGAGGATCCTTTCTACAAAGTAAAGGAGGTATTACCTAAGGAACTTACTGGTTTACGTATT<br>CGAGTAAACAAGGCTGTCATGAATGCAACAAAGGGATTGTCTAAGGATAAATTCAATTACGGG<br>CCTCATGACTTCAGCTTAGCAGCCAGAAATGGAATTTGTTTTGCGTTCAGAGAGGTCGAC |
| pSC3718 | TCTAGAACCGAACAGACGACCCATTTCCTGTGGCAGGGTTATCGGTTGCTGCAGGAGCAGCGC<br>GACGACGGCAGCCGCCGCAGCTGGAGCTACGATCCGGCCAGCCCGTGGAGCCCGTTGGCGGCG<br>CTGGAGCAGGCAGGCGACAGCCGCTCGGCGGATATTTACTGGTATCACACCGATCTGAACAGC<br>GCGCCGCTGGAGGTGACCGACGCGGCGGGCAACCTGTGCTGGTCCGGGCAATACGACACCTTC<br>GGCAAGCTGCAGGGCCAGACGGTGGCCGGCGCGGCGAAGCGGCAGGGCGCGCAATACCAGCAG<br>CCGCTGCGCTACGCCGGGCAATACCAGGATGACGAAAGTGGCCTGCACTACAACCTGTTCCGC<br>TACTACGAACCCGAGGTGGGGCGTTTCACCACGCAGGATCCGATAGGGTTGGAAGGCGGGCTG<br>AATTTATATGCTTATGGGCCAAATCCTTTAACTTGGATAGATCCATTTGGGTTAAGTAATTGT<br>TCGACTCTTGACCGTATTATTGGTGATGCGAACAAAGTCGCTTCGCGTGGTGGAGCTATAACA<br>GCAAAACAAGCTCAGATACTGAGAGATAATTTACCGGTAGTTCAGAGACGGAGTGTTTTCCAA<br>AATCAGATGGCTCGCAAACGGTTCGTCGCAGATCAGCATTATCTGATGAGTCAGTGGGAAGCC<br>AATACAGGTAGAACCTGGCCGACAGGAGCTACCCCGCACCACATAATCCCACTAGAAAGTGGG<br>GGGTGGAATGAATGGTGGAATTTGATGCCTACTCATGGCACTTTGCCTAATCATTCTTTACCT<br>GGTGTACCTGGCCCACATGCCGCCGGTGGAGTGCTTCGAACAACTGTTCAACAAAGCAGAAAA<br>GCACTGCCTCCTGGAACTATAACTGATTTGAGATTGTGACATGAATGAATTTGACTTTGATAG<br>TTTACTACAGCGCATCGATAGCTCATGCTTTTTCTCAAGAATGGGGCTCCCAGATGTTCTGGA<br>TAGTCGCGTGATTTTAATTGAAAATGTTGAGAAGGTGTTTGTTAATCCCACTGATGCAGAGTT<br>TAAGGGATACTATGATAGTGTGGAATGGTTGCCAACATCAATGACTCAGGAGGATCCTTTCTA<br>CAAAGTAAAGGAGGTATTACCTAAGGAACTTACTGGTTTACGTATTCGAGTAAACAAGGCTGT<br>CATGAATGCAACAAAGGGATTGTCTAAGGATAAATTCAATTACGGGCCTCATGACTTCAGCTT<br>AGCAGCCAGAAATGGAATTTGTTTTGCGTTCAGAGAGTATGTTTCTGAACAGTATCTTCATTT<br>AGGGAATAAGTGGGAAGAAGTTGTGGGTATATACTTTTCTGGTCACTGGCCTGTGGGTATAGC<br>AAAAGATAAGATAGTTACCATTTAAGGGCCC |

**Supplementary Table 3. Crystallographic statistics.**

| | Rhs2CT-RhsI2-EF-Tu | Rhs2CT-RhsI2 |
|---|---|---|
| Wavelength (Å) | 0.97628 | 0.87313 |
| Space group | *P* 1 21 1 | *P* 21 21 21 |
| Beamline | I03 | ID23-1 |
| Subunits/asymmetric unit | 2 | 1 |
| Unit cell dimensions *a*, *b*, *c* (Å) | 61.60 110.57 113.86 | 37.01 81.93 101.37 |
| Unit cell angles *α*, *β*, *γ* (º) | 90.00 102.39 90.00 | 90.00 90.0 90.00 |
| Resolution range (Å) | 60.24 - 2.45 | 2.17 - 37.98 |
| Total number of reflections | 345063 | 102443 |
| Unique reflections | 51653 | 17007 |
| Completeness (%)[a] | 100 (100.0) | 100.0 (99.2) |
| $R_{merge}$[b] | 0.157 (1.444) | 0.157 (1.511) |
| Redundancy | 6.7 (6.1) | 6.0 (6.2) |
| $R_{pim}$[c] | 0.067 (0.636) | 0.07 (0.661) |
| $\langle I/\sigma(I)\rangle$ | 6.9 (1.0) | 8.7 (3.0) |
| Wilson *B*-factor (Å$^2$) | 46.5 | 26.0 |
| CC(1/2) | 0.988 (0.604) | 0.993 (0.542) |
| **Refinement** | | |
| $R_{work}$[d]/$R_{free}$[e] (%) | 0.27 / 0.31 | 0.19 / 0.25 |
| Number of reflections for $R_{work}/R_{free}$ | 52057 / 2724 | 16107 / 842 |
| Number of protein residues | 1313 | 272 |
| Number of GDP / IMD / EDO / $Mg^{2+}$ | 2 / 2 / 2 / 2 | NA |
| Number of $CH_3COO^-$ / $C_2H_6O_2$ | NA | 3/1 |
| DPI[f] (Å) | 0.34 | 0.20 |
| RMSD Bond lengths (Å)/Angles (º) | 0.0063 / 1.2801 | 0.007 / 1.48 |
| Average *B*-factors (Å$^2$) per chain | 52.84 / 63.44 / 62.65 / 52.76 / 62.1 / 62.2 / 52.0 | 35.50 / 37.48 |
| Water molecules (Average *B*-factors, Å$^2$) | 146 (41.44) | 131 (39.89) |
| Ligands (Average B-factors, Å$^2$) | GDP (39.8) Imidazole (56.9) $C_2H_6O_2$ (54.8) $Mg^{2+}$ (31.0) | $CH_3COO^-$ (47.5) $C_2H_6O_2$ (50.4) |
| **Ramachandran analysis** | | |
| Residues in favoured regions (%) | 94.4 | 95.9 |
| Residues in allowed regions (%) | 5.5 | 3.73 |
| Rotamer outliers (%) | 0.15 | 0.37 |

[a.] The default spherical completeness is reported. [b.] $R_{merge} = \sum h \sum i |I[h,i] - \langle I[h]\rangle| / \sum h \sum i\, I[h,i]$; where $I[h,i]$ is the intensity of the *i*th measurement of reflection *h* and $\langle I[h]\rangle$ is the mean value of $I[h,i]$ for all *i* measurements. [c.] $R_{pim} = \sum h\, [1/[/nh\text{-}1]]^{1/2} \sum i |\langle Ih\rangle\text{-}\, Ih,i| / \sum h \sum i\, Ih,i$. [d.] $R_{work} = \sum hkl ||F_o|\text{-}|F_c|| / \sum |F_o|$, where $F_o$ is the observed structure factor amplitude and $F_c$ is the structure factor amplitude calculated from the model. [e.] $R_{free}$ is calculated with a subset of data that are excluded from refinement calculations (5%) using the same method as for $R_{merge}$. [f.] DPI = Diffraction-component Precision Index as defined by Cruickshank[15].

### Supplementary Table 4. Protein purification details.

| Plasmid name | Plasmid encodes | Purified protein or complex | Extinction coefficent $M^{-1}$ $cm^{-1}$ at 280 nm | Purification step | Buffer |
|---|---|---|---|---|---|
| pSC3502 | $His_6$-Rhs2CT RhsI2 | Rhs2CT-RhsI2-EF-Tu complex | 70820 | HisTrap HP 5 mL $Ni^{2+}$-affinity column | Buffer *A1*: 50 mM MES-HCl 50 mM imidazole-HCl pH5.5, 250 mM NaCl, 1 mM $MgCl_2$ and 0.5 mM TCEP (500 mM imidazole in buffer *B1*) |
| | | | | Reverse purification HisTrap HP 5 mL $Ni^{2+}$-affinity column | Buffer *A1*: 50 mM MES-HCl 50 mM imidazole-HCl pH5.5, 250 mM NaCl, 1 mM $MgCl_2$ and 0.5 mM TCEP (500 mM imidazole in buffer *B1*) |
| | | | | Superdex 75 HiLoad 26/60 column | Buffer *C1*: 25 mM MES-HCl 25 mM imidazole-HCl pH5.5, 150 mM NaCl, 1 mM $MgCl_2$ and 0.5 mM TCEP |
| | | $His_6$-Rhs2CT (refolded) | 23490 | HisTrap HP 5 mL $Ni^{2+}$-affinity column | Buffer *A2*: 50 mM Tris-HCl pH 7.5, 300 mM NaCl, 0.5 mM TCEP and 20 µM $Zn(OAc)_2$ (500 mM imidazole in buffer *B2*) |
| | | | | Superdex 75 HiLoad 16/60 column | Buffer *C2*: 25 mM Tris-HCl pH 7.5, 150 mM NaCl, 0.5 mM TCEP and 20 µM $Zn(OAc)_2$ |
| | | $His_6$-Rhs2CT (refolded)-$His_6$-RhsI2 complex | 50420 | Superdex 75 HiLoad 16/60 column | Buffer *C3*: 20 mM MES-HCl 20 mM imidazole-HCl pH6.0, 150 mM NaCl, 1 mM $MgCl_2$ and 0.5 mM TCEP |
| pSC3251 | $His_6$-$Rhs2CT_{CM}$ | $Rhs2CT_{CM}$-EF-Tu complex | 43890 | HisTrap HP 5 mL $Ni^{2+}$-affinity column | Buffer *A3*: 50 mM Tris-HCl pH 8.0, 250 mM NaCl, 0.5 mM TCEP and 1 mM $MgCl_2$ (500 mM imidazole in buffer *B3*) |
| | | | | Reverse purification HisTrap HP 5 mL $Ni^{2+}$-affinity column | Buffer *A3*: 50 mM Tris-HCl pH 8.0, 250 mM NaCl, 0.5 mM TCEP and 1 mM $MgCl_2$ (500 mM imidazole in buffer *B3*) |
| | | | | Superdex 75 HiLoad 16/60 column | Buffer *C3*: 50 mM Tris-HCl pH 8.0, 250 mM NaCl, 0.5 mM TCEP and 1 mM $MgCl_2$ |
| pSC3712 | $His_6$-$Rhs2CT_{EM\text{-}CM}$ RhsI2 | $Rhs2CT_{EM\text{-}CM}$-RhsI2 complex | 55920 | HisTrap HP 5 mL $Ni^{2+}$-affinity column | Buffer *A4*: 50 mM Tris-HCl pH 7.3, 250 mM NaCl, 20 mM imidazole, 1 mM $MgCl_2$, 0.5 mM TCEP and 30 µM $Zn(OAc)_2$ (500 mM imidazole in buffer *B4*) |
| | | | | Reverse purification HisTrap HP 5 mL $Ni^{2+}$-affinity column | Buffer *A4*: 50 mM Tris-HCl, pH 7.3, 250 mM NaCl, 20 mM imidazole, 1 mM $MgCl_2$, 0.5 mM TCEP and 30 µM $Zn(OAc)_2$ (500 mM imidazole in buffer *B4*) |
| | | | | Superdex 75 HiLoad 16/60 column | Buffer *C4*: 25 mM MES-HCl 25 mM imidazole-HCl pH5.5, 150 mM NaCl, 1 mM $MgCl_2$ and 0.5 mM TCEP |
| | | $Rhs2CT_{EM\text{-}CM}$ (refolded) | 28990 | HisTrap HP 5 mL $Ni^{2+}$-affinity column | Buffer *A1*: 50 mM MES-HCl 50 mM imidazole-HCl pH5.5, 250 mM NaCl, 1 mM $MgCl_2$ and 0.5 mM TCEP (500 mM imidazole in buffer *B1*) |
| | | | | Superdex 75 HiLoad 16/60 column | Buffer *C1*: 25 mM MES-HCl 25 mM imidazole-HCl pH5.5, 150 mM NaCl, 1 mM $MgCl_2$ and 0.5 mM TCEP |
| pSC955 | $His_6$-RhsI2 | $His_6$-RhsI2 | 26930 | HisTrap HP 5 mL $Ni^{2+}$-affinity column | Buffer *A2*: 50 mM Tris-HCl pH 7.5, 250 mM NaCl (500 mM imidazole in buffer *B2*) |
